# Resolving the graft ischemia-reperfusion injury during liver transplantation at the single cell resolution

**DOI:** 10.1101/2021.01.10.426087

**Authors:** Linhe Wang, Jie Li, Shuai He, Haitian Chen, Shujiao He, Meixian Yin, DaWei Zou, Jinghong Xu, Shirui Chen, Tao Luo, Xinyu Yu, Jinxin Bei, Zhiyong Guo, Xiaoshun He

**Author notes:** Correspondence should be addressed to (JXB), or (ZYG), or (XSH). Email addresses of authors: Linhe Wang,; Jie Li; Shuai He,; Haitian Chen; Shujiao He; Meixian Yin; DaWei Zou; Jinghong Xu,; Shirui Chen; Tao Luo; Xinyu Yu; Jinxin Bei,; Zhiyong Guo,; Xiaoshun He. These authors contributed equally (Linhe Wang, Jie Li, Shuai He).

## Abstract

Ischemia-reperfusion injury (IRI) remains the major reason for impaired donor graft function and increased mortality post-liver transplantation. The mechanism of IRI involves multiple pathophysiological processes and numerous types of cells. However, a systematic and comprehensive single-cell transcriptional profile of intrahepatic cells during liver transplantation is still unclear. We performed a single-cell transcriptome analysis of 14,313 cells from liver tissues collected from pre-procurement, at the end of preservation and 2 h post-reperfusion. We made detailed annotations of mononuclear phagocyte, endothelial cell, NK/T, B and plasma cell clusters, and we described the dynamic changes of the transcriptome of these clusters during IRI and the interaction between mononuclear phagocyte clusters and other cell clusters. In addition, we found that TNFAIP3 interacting protein 3 (*TNIP3)*, specifically and highly expressed in Kupffer cell clusters post-reperfusion, may have a protective effect on IRI. In summary, our study provides the first dynamic transcriptome map of intrahepatic cell clusters during liver transplantation at single-cell resolution.

## 1 INTRODUCTION

Liver transplantation is the standard therapy for end-stage liver disease ^1^. At present, almost all transplanted livers suffer ischemia during cold preservation and subsequent reperfusion injury ^2^, posing a huge challenge to the functional recovery of donor livers and the prognosis of recipients. The mechanism involved in ischemia-reperfusion injury (IRI) is complex, including changes in a variety of cellular components, inflammatory factors and mediators ^3-5^. For example, hepatocytes and liver sinusoidal endothelial cells (LSEC) are sensitive to ischemia. The lack of oxygen leads to disorders of the respiratory chain, accelerated glycolysis, and electrolyte disturbances, which in turn leads to microcirculation disorders and impaired cell functions ^6,7^. After reperfusion, harmful molecules especially lipopolysaccharide (LPS) enters the liver and activates Kupffer cells, produces reactive free radicals (ROS) and pro-inflammatory cytokines, including tumor necrosis factor-α (TNF-α) and interleukin-1β (IL-1β), which triggers the inflammatory cascade and cell apoptosis after IRI ^8,9^.

Immune cells and non-immune cells in the liver are heterogeneous and consist of multiple subpopulations of various immunological and physiological functions, including T cells, nature killer (NK) cells, B cells, plasma cells, mononuclear phagocytes and endothelial cells ^10^. The flow cytometry used in previous studies can only target specific cell subpopulations and marker genes ^11-13^. The recent development of unbiased single-cell RNA sequencing (scRNA-seq) technology can well identify different cell populations and explain the heterogeneity and relevance between cells. The scRNA-seq has annotated the cellular transcription profile of adult liver tissues ^14^ and revealed the molecular mechanism of myocardial ischemia injury ^15,16^. However, the cellular and molecular mechanism of graft IRI during liver transplantation is still unknown at the single-cell resolution level.

Herein, we used scRNA-seq to obtain the first unbiased and comprehensive liver transplant cell atlas by collecting liver tissue samples pre-procurement (PP), at the end of preservation (EP) and 2 h post-reperfusion (PR). This atlas annotated different cell subgroups, revealed their changes in the transcriptome and illustrated the interactions between different cell subpopulations during liver transplantation. This research will serve as an important resource to further understand the cellular and molecular mechanism of graft IRI during liver transplantation.

## 2 RESULTS

### 2.1 ScRNA-seq identifies intrahepatic cell subpopulations

We performed scRNA-seq of intrahepatic cells in PP, EP and PR samples from an adult donor liver. A total of 18,793 single-cell transcriptomes were obtained using the 10X Genomics platform (**Figure 1A**). We identified 1160 doublets using “DoubletFinder” software (Figure S1A, Supporting Information). Cells with low quality (UMI < 1,000, gene number < 500) and high (> 0.25) mitochondrial genome transcript ratio were removed. Finally, a total of 14,313 single-cell transcriptomes were obtained for the further analysis. Changes in cell type-specific gene expression caused by the IRI response can make joint analysis difficult. Therefore, we performed an integrated analysis for cell type identification and comparison using Seurat R package v3.

**Figure 1:**
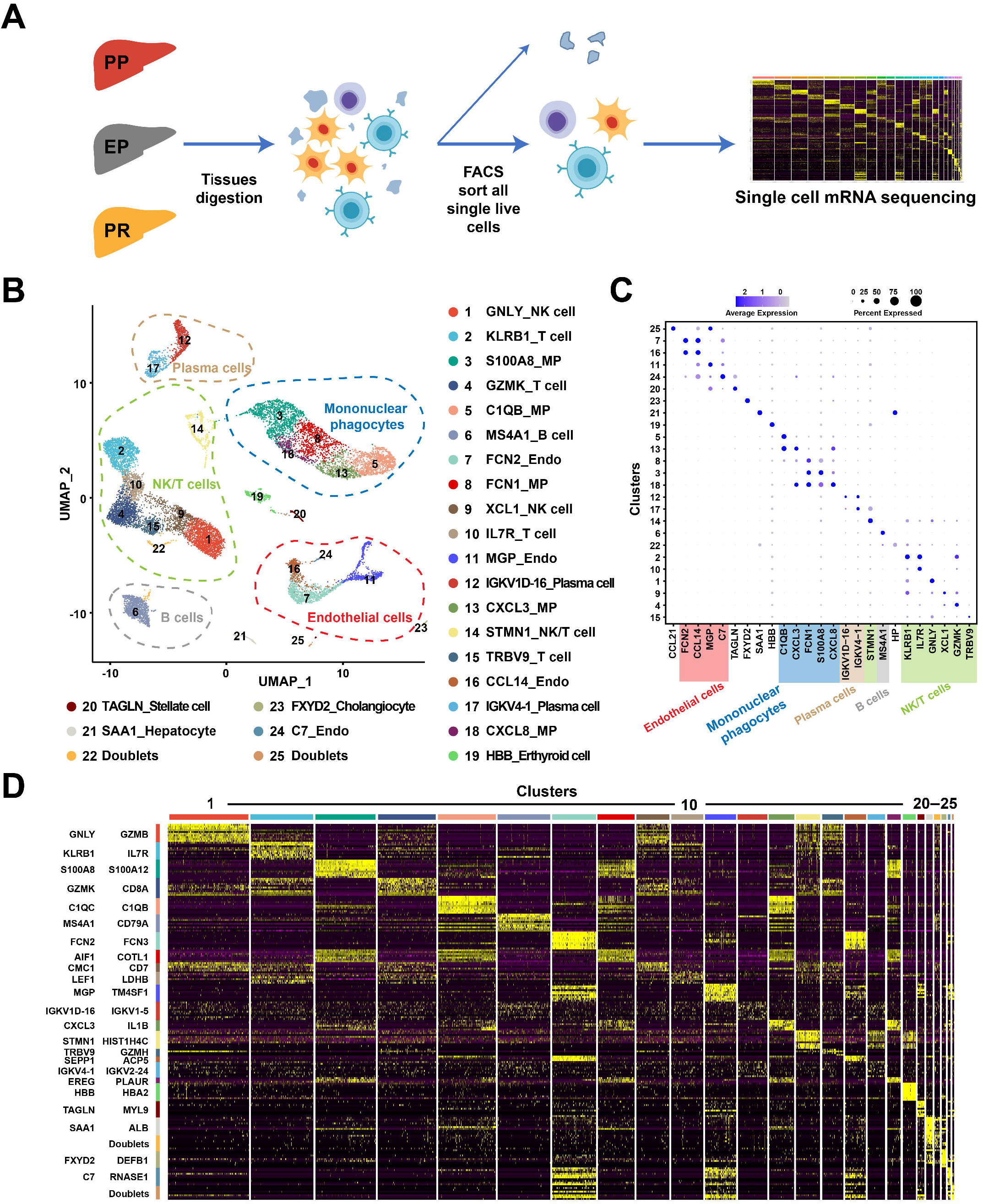
Overview of scRNA-seq from samples in liver transplantation. **A**: Workflow of tissue collection, sample processing and data acquisition. Collect samples from three time points, PP, EP and PR, for tissue digestion. Next, sort live cells by FACS and construct cDNA libraries, then perform high-throughput sequencing and downstream analyses. **B**: UMAP visualization of all cells (14,313) in 25 clusters. Each dot represents one cell, with colors coded according to the different cluster. Clusters are named by the most specific and highly expressed genes. MP: mononuclear phagocytes; Endo: endothelial cells. **C**: Dot plots showing the most highly expressed marker genes (x axis) of major cell types (y axis) in **Figure 1B**. The color of the dots represents the level of gene expression while the size of the dot represents the percent of cells expressing the gene. **D**: Heatmap of top 10 differentially expressed genes between different clusters. The line is colored according to clusters in **Figure 1B**. Each cluster lists the top two genes shown on the left.

Clustering the intrahepatic cells revealed 25 populations (Figure 1B). We named these clusters with its most specifically and highly expressed marker genes (Figure 1C). And the differential gene expression (DEG) analysis showed that each cluster had a specific gene signature (Figure 1D). These clusters across five major cell lineages including mononuclear phagocytes (MP), endothelial cells, NK/T, B and plasma cells, were identified by their canonical signature gene profiles such as *CD68* (macrophage marker), *CD3D* (T cell marker), *FCGR3A* (NK cell marker), *MS4A1* (B cell marker), *SDC1* (plasma cell marker), and *PECAM1* (endothelial marker), respectively (Figure 1B, C, Figure S1B, Table S1, Supporting Information). These five populations contained cells from samples collected at all of three time points (PP, EP and PR) (Figure S1C, Supporting Information).

### 2.2 Different pro- and anti-inflammatory phenotypes in mononuclear phagocytes

We obtained 3,622 mononuclear phagocytes from PP, EP and PR samples, which were re-grouped into four distinct clusters (**Figure 2A** left panel, Figure S2A, Supporting Information), annotated as two tissue monocytes (with suffix of TMo, highly expressed *S100A9* gene) and two Kupffer cell clusters (with suffix of KC, highly expressed *C1QC* gene) (Figure 2B), based on the most specific, highly expressed gene in that cluster. The hierarchical clustering analysis revealed that the two KC clusters were closely related as a branch node, as were the two TMo clusters (Figure 2A right panel).

**Figure 2:**
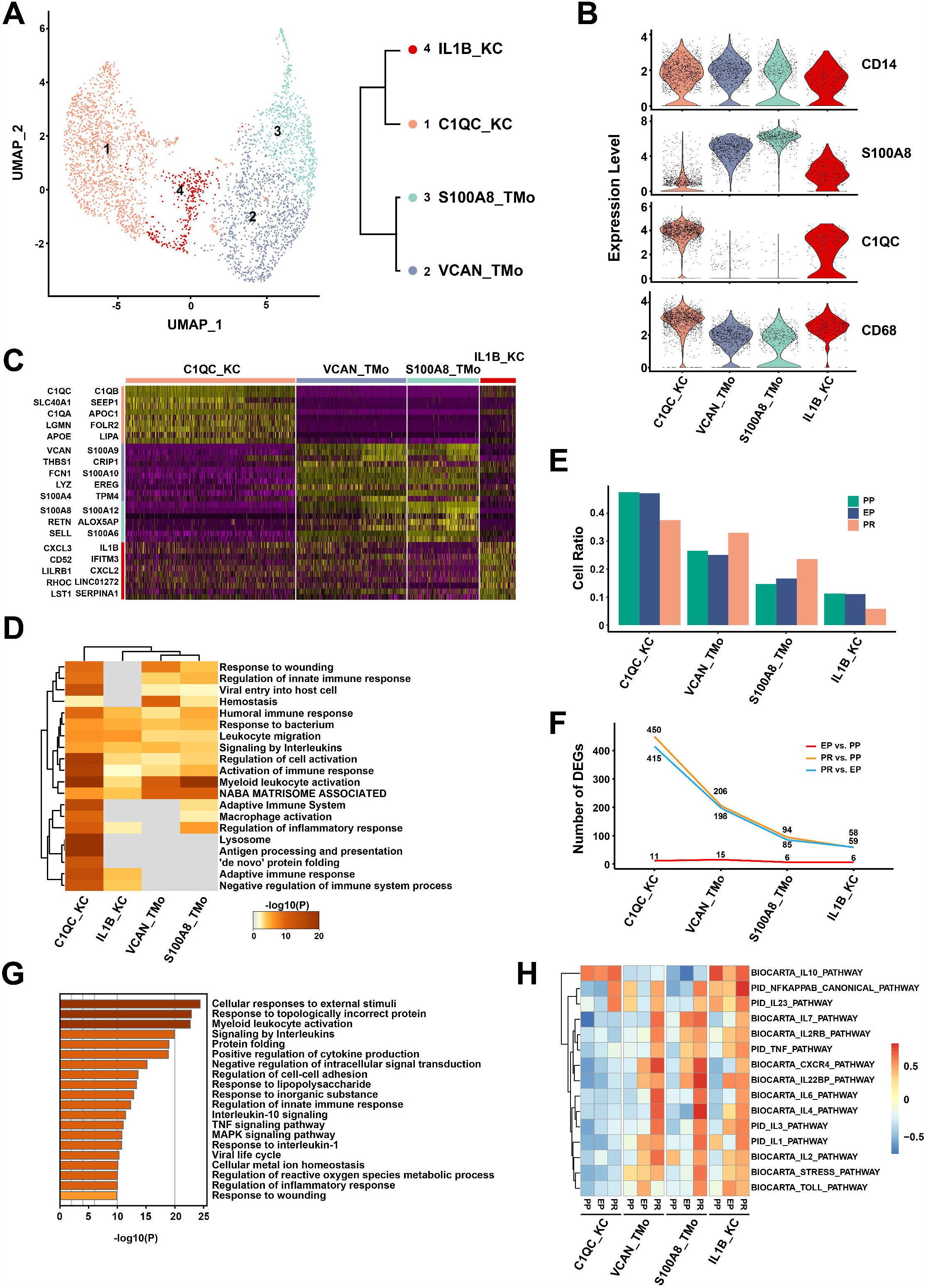
scRNA-seq of mononuclear phagocytes in liver transplantation. **A**: UMAP plot showing four mononuclear phagocyte clusters (3622 cells) in liver transplantation, colored according to different clusters (left panel). Dendrogram of four clusters by the hierarchical clustering analysis based on their normalized mean expression values (right panel). **B**: Violin plots showing the normalized expression of *CD14, S100A8, C1QC* and *CD68* genes (y axis) for TMo and KC clusters (x axis). **C**: Heatmap of top 10 differentially expressed genes between different mononuclear phagocyte clusters. The line is colored according to clusters in **Figure 2A**. **D**: Gene Ontology enrichment analysis results of mononuclear phagocyte clusters. Only the top 20 most significant GO terms (*P*-value < 0.05) are shown in rows. **E**: Cell ratio of different mononuclear phagocyte clusters in PP, EP and PR samples. **F**: Number of up-regulated DEGs between different timepoint samples in different mononuclear phagocyte clusters. **G**: Gene Ontology enrichment analysis results of up-regulated DEGs in C1QC_KC cluster after reperfusion (PR vs. EP). Only the top 20 significant GO terms (*P*-value < 0.05) are shown in rows. **H**: The gene set variation analysis (GSVA) showing the pathways (PID and BIOCARTA gene sets) with significantly different activation in different samples of mononuclear phagocyte clusters. Different colors represent different activation scores.

VCAN_TMo and S100A8_TMo clusters highly expressed *S100A12, S100A8* and other genes similar to those expressed by peripheral monocytes, indicating that cells in these two clusters might just be recruited and differentiated from peripheral blood circulating monocytes. In VCAN_TMo cluster, the five most highly expressed genes were *VCAN, S100A9, THBS1, CRIP1* and *FCN1* while in S100A8_TMo cluster were *S100A8, S100A12, RETN, ALOX5AP* and *S100A9*. C1QC_KC and IL1B_KC clusters highly expressed *C1QC, C1QB* and other tissue resident marker genes in macrophages, indicating that they are Kupffer cells, and the top five highly expressed genes were *C1QC, C1QB, SLC40A1, SEPP1, C1QA* in C1QC_KC and *IL1B, CXCL3, IFITM3, CD52, CXCL2* in IL1B_KC (Figure 2C).

We investigated the potential biological functions of different TMo and KC clusters using gene ontology (GO) analysis with up-regulated DEGs, and the analysis results revealed the differences and commonalities between the various clusters (Table S2, Supporting Information). The biological function commonly enriched by TMo and KC clusters were *Myeloid leukocyte activation, Leukocyte migration, Signaling by interleukins* and other pathways of immune response (Figure 2D). In addition, C1QC_KC cluster showed strong immune regulatory ability and high enrichment in *Regulation of protein stability, Regulation of tumor necrosis factor production, Antigen processing and presentation* and other pathways. Most of the pathways enriched in IL1B_KC cluster were the same as those of in C1QC_KC cluster, except pathways such as *Negative regulation of viral process*. S100A8_TMo cluster was enriched in *Formyl peptide receptors bind formyl peptides* and *Defense response to fungus* pathways except for the common enrichment immune response related pathways, while VCAN_TMo cluster was additionally enriched in *Dissolution of Fibrin Clot, Positive regulation of organelle organization* and other pathways (Figure S2B, Supporting Information).

In order to explore the dynamic changes of different TMo and KC clusters during liver transplantation, we performed a comparative analysis between EP and PP samples in the cold preservation stage (EP vs. PP), PR and EP samples in the reperfusion stage (PR vs. EP), as well as PR and PP samples in the overall stage (PR vs. PP). In the PP and EP samples, we found that the proportion of each cluster in the corresponding sample did not change significantly. In the PR sample, the proportion of KC clusters was lower than that of PP and EP samples, while the proportion of TMo clusters was higher than those of PP and EP samples (Figure 2E).

Next, we performed DEG analysis and found that only a small number of differentially expressed genes were up-regulated in each cluster during the cold preservation stage. In contrast, more genes were up-regulated in each cluster during reperfusion stage and overall stage when compared with the cold preservation stage respectively. The number of up-regulated genes between reperfusion stage and overall stage was very close, but the amount of DEG up-regulation in the overall stage was slightly more than that in reperfusion stage. These two stages shared most of the genes that were up-regulated, especially those with high expression levels; a similar trend was observed in the down-regulated genes (Figure 2F, Figure S2C-D, Table S3-5, Supporting Information).

The results of GO analysis showed that C1QC_KC cluster up-regulated multiple stress, inflammatory response pathways such as *Cellular responses to external stimuli, Response to topologically incorrect protein* and cell activation pathways such as *Myeloid leukocyte activation* in the reperfusion stage (Figure 2G). Like C1QC_KC cluster, the IL1B_KC, VCAN_TMo and S100A8_TMo clusters also up-regulated stress response pathways and cell activation pathways in the reperfusion stage (Figure S3A left panel, Supporting Information). The most obvious down-regulated pathways for C1QC_KC, IL1B_KC, VCAN_TMo and S100A8_TMo clusters in the reperfusion stage are *G protein-coupled purinergic receptor signaling pathway, Influenza A, Response to antibiotic, Response to inorganic substance* respectively. (Figure S3A right panel, Supporting Information). This phenomenon was similarly found in the overall stage with the reperfusion stage, there was no obvious enrichment pathway in the cold preservation stage because of the small number of DEGs (Figure S3B, Supporting Information).

In order to further explore the changes of specific inflammation-related pathways in different stages, we performed gene set variation analysis (GSVA) and selected the pathways related to IRI. The results showed that various pro-inflammatory pathways such as *IL1 pathway, IL2 pathway, TNF pathway* and *NFKAPPAB pathway* were activated after reperfusion in VCAN_TMo, S100A8_TMo and IL1B_KC clusters, while these pathways maintained a low activation level in C1QC_KC cluster during the cold preservation and reperfusion stage. In contrast, the *IL10 pathway*, which is considered as protective during IRI ^17^, were significantly activated in C1QC_KC and IL1B_KC cluster of samples collected at various timepoints. This result suggests that C1QC_KC cluster may have an anti-inflammatory regulatory effect in IRI, while VCAN_TMo and S100A8_TMo clusters may play a pro-inflammatory role in IRI. IL1B_KC cluster tends to be an intermediate cell population, which has both pro-inflammatory and anti-inflammatory effects (Figure 2H). In addition, we assessed the antigen-presenting abilities for extracellular antigens as reflected by antigen-presenting score (APS), and found that the KC clusters, particularly the C1QC_KC cluster, had higher APS than the TMo clusters in MHC class II genes (*P*<0.0001) (Figure S2E, Supporting Information).

### 2.3 Specific up-regulation of TNIP3 in KC clusters after reperfusion

To further explore the anti-inflammatory mechanism of KC clusters, we studied the DEGs that may be related to the anti-inflammatory effect during the cold preservation stage and the reperfusion stage. In the reperfusion stage, in addition to the heat shock protein family proteins and metal ion binding proteins, the role of which are already well known in IRI, we found that nuclear factor kappa-B (NF-κB) activation inhibitor *TNIP3* was specifically and highly expressed in the KC clusters after reperfusion (Fold Change (FC)=2.2 in PR vs. EP of C1QC_KC cluster; FC=3 in PR vs. EP of IL1B_KC cluster).

*TNIP3* was almost not expressed in PP and EP samples, while the expression level in PR sample was significantly increased. Interestingly, it was almost only expressed in the KC clusters especially in the C1QC_KC cluster when compared with other intrahepatic cells in our dataset (**Figure 3A, B**). In addition, the high expression of *TNIP3* after reperfusion (log_2_ FC=3.91) was verified in our bulk RNA sequencing data from 14 EP-PR pairs of donor liver samples (Table S6, Supporting Information). Immunofluorescence results showed that in other four paired samples, CD68+ cells did not express TNIP3 in the EP samples, while a proportion of CD68+ cells clearly expressed TNIP3 protein in the PR samples (Figure 3C). The Western Blot results also showed that the expression of TNIP3 protein in the PR group was higher than that in the PP and EP group (Figure 3D).

**Figure 3:**
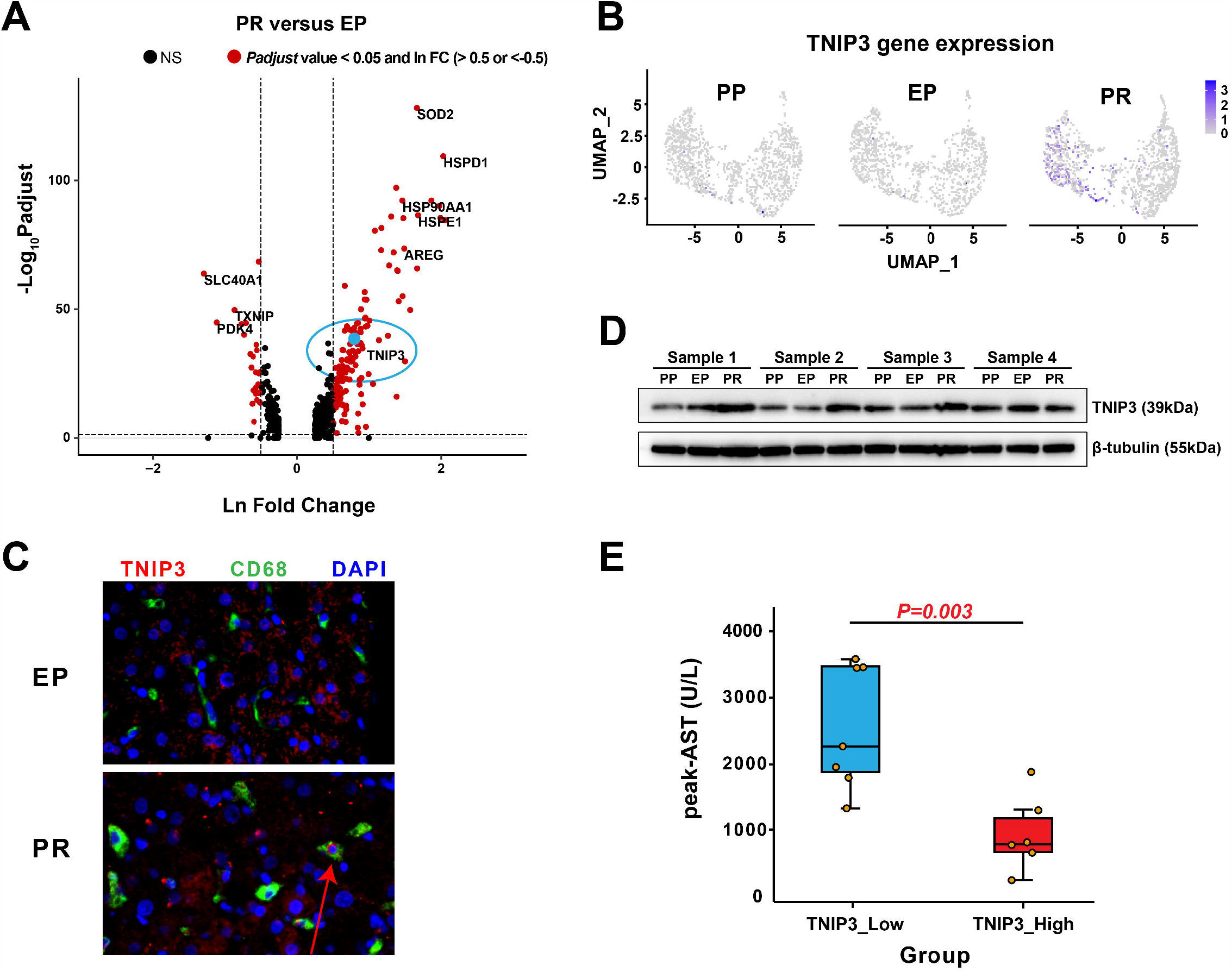
TNIP3 expression is specifically increased in KC clusters after IRI. **A**: Volcano plot shows the DEGs of PR vs. EP in C1QC_KC cluster. The red dots represent *P*-adjust value less than 0.05 and lnFC greater than 0.5 or less than −0.5, the blue dot represent *TNIP3*, and the rest are black dots. **B**: Feature plots showing the normalized expression of *TNIP3* in mononuclear phagocyte clusters of PP, EP and PR. Color represents the level of gene expression. **C**: Immunofluorescence showed that the expression of TNIP3 (red), CD68 (green) and DAPI (purple) in EP and PR samples. CD68+TNIP3+ cells only appeared in partly cells in PR sample. **D**: Western Blot results showed that the expression of TNIP3 in PR samples was higher than that in the EP and PP samples. **E**: The postoperative peak-AST level in the TNIP3_low group is higher than TNIP3_high group (n=13, 2506 vs. 880 u/l, *P*=0.003), each point represents a patient.

To further explore the role of *TNIP3* in IRI, the expression level of this gene in 14 PR samples was analyzed and the patients were divided into *TNIP3* high expression group and low expression group according to the median value of FC (3.438). We found that the peak aspartate aminotransferase (AST) levels of patients within 7 days post-transplantation in the high expression group were significantly lower than those in low expression group (880 vs. 2506 U/L, *P*=0.003) (Figure 3E, Table S7, Supporting Information), suggesting a protective role of *TNIP3* in graft IRI.

### 2.4 Endothelial cells in liver transplant IRI

We obtained single cell transcriptomes of 1,766 cells from the endothelial cell lineage of three samples and re-clustered them into seven clusters (**Figure 4A**, Figure S4A, Supporting Information). These clusters uncovered two major populations of endothelial cells in the liver with different canonical marker genes, the liver sinusoidal endothelial cells (LSEC) and vascular endothelial cells (VEC). LSEC is a group of endothelial cells with different functions and morphologies that exist in the liver sinusoids with filtering and scavenging roles ^18,19^. The VEC mainly lined in the hepatic artery and vein ^20^. We annotated the four *CLEC4G*^*+*^*PECAM1*^*low*^ clusters as LSEC based on previous studies ^14,21-23^. Their highly expressed top five genes were *CTSL, CLEC4M, CCL23, CD14*, and *FCGR2B* in CTSL_LSEC; *BGN, CPM, IGFBP3, CD24*, and *PLPP3* in BGN_LSEC; *STAB1, MT1G, MEG3, SERPINA1*, and *NEAT1* in STAB1_LSEC; *NUPR1, GPX4, CSTB, PVALB* and *POLR2L* in NUPR1_LSEC.

**Figure 4:**
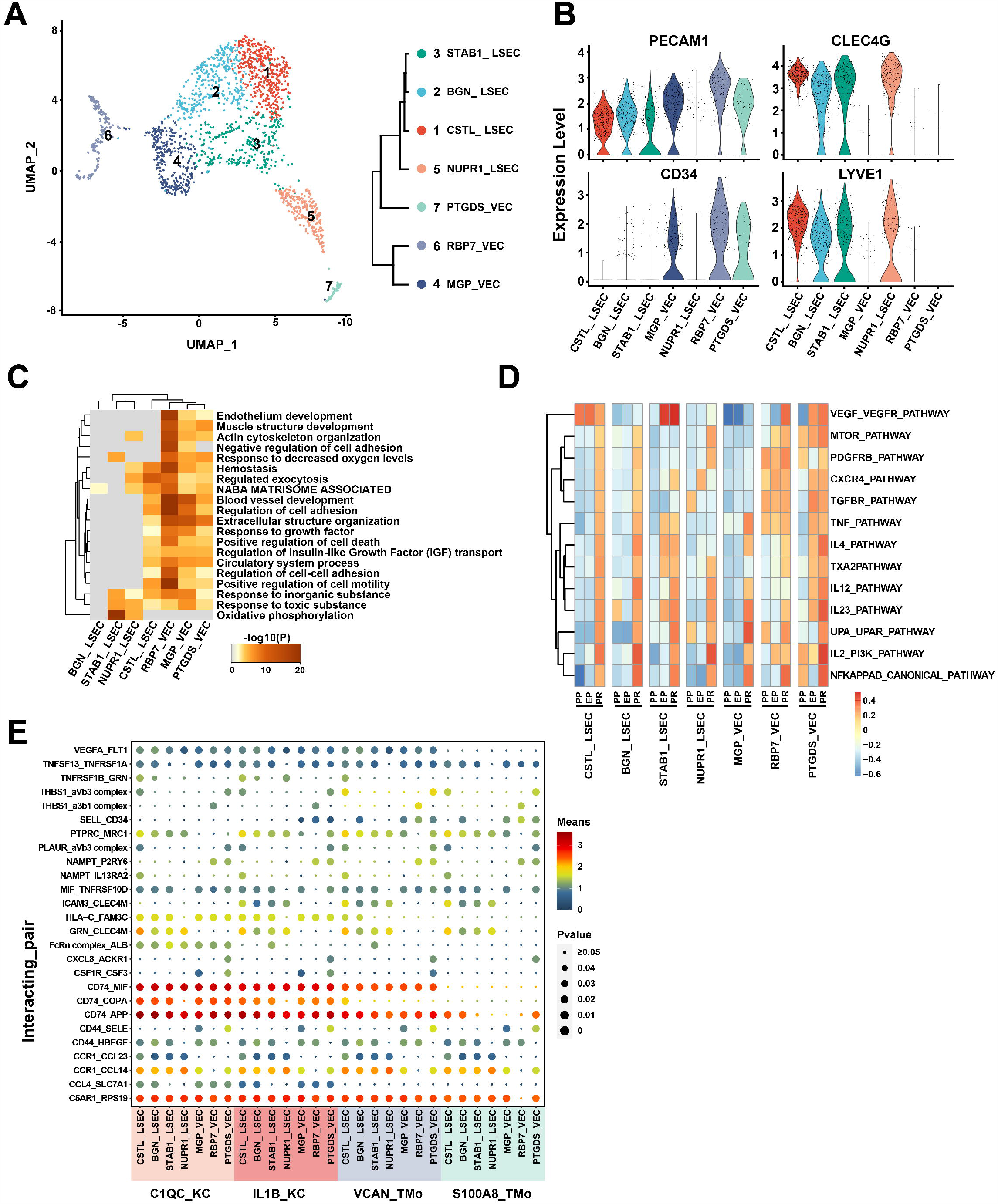
scRNA-seq of endothelial cells in liver transplantation. **A**: UMAP plot showing seven endothelial cell clusters in liver transplantation, colored according to different clusters (left panel). Dendrogram of seven clusters by hierarchical clustering analysis based on their normalized mean expression values (right panel). **B**: Violin plots showing the normalized expression of *PECAM1, CLEC4G, CD34* and *LYVE1* genes (y axis) for endothelial cell clusters (x axis). **C**: Gene Ontology enrichment analysis results of endothelial cell clusters. Only the top 20 most significant GO terms (*P*-value < 0.05) are shown in rows. **D**: GSVA showing the pathways (PID gene sets) with significantly different activation in different samples of endothelial cell clusters. Different colors represent different activation scores. **E**: Cell-cell interaction analysis between mononuclear phagocyte clusters and different endothelial cell clusters in PR samples. Ligand-receptor pairs are labeled in y-axis. The size of the circle represents the level of p-value while different colors represent different means value. Ligands come from mononuclear phagocyte clusters, and receptors come from endothelial cell clusters.

The other three clusters were *CD34*^*+*^*PECAM1*^*high*^ and annotated as VEC based on published literatures ^14,21-23^. Their top five highly expressed genes were *MGP, ADIRF, CLU, MT1M*, and *CPE* in MGP_VEC; *RBP7, GSN, RGCC, CXCL12*, and *PLVAP* in RBP7_VEC; *PTGDS, ADGRG6, POSTN, TAGLN*, and *IL1RL1* in PTGDS_VEC (Figure 4B, Figure S4B, Table S8, Supporting Information).

In order to compare the differences in the biological functions of different endothelial cell clusters, we performed GO analysis in the seven endothelial cell clusters. The VEC clusters were enriched in *Extracellular structure organization, Muscle structure development, Endothelium development* and other pathways while the LSEC clusters were quite different. STAB1_LSEC and NUPR1_LSEC clusters were specifically enriched in *Oxidative phosphorylation* pathway. Many pathways enriched in CSTL_LSEC were similar to those of VEC clusters. There were few DEG enrichment pathways in BGN_LSEC cluster (Figure 4C).

In the study of the dynamic changes of various endothelial cell clusters in liver transplantation, we found that there were few genes differentially expressed in each cluster during the cold preservation stage, while the number of DEGs significantly increased in the reperfusion stage and the overall stage, and most of the high expression level genes are shared between them (Figure S4C-D, Table S9-11, Supporting Information). The GO analysis of DEGs showed that almost all endothelial cell clusters had up-regulation of *Protein folding* or *Response to topologically incorrect protein* related to stress response and cell adhesion related inflammation-regulation pathways in the reperfusion stage. In addition, the CSTL_LSEC and BGN_LSEC clusters also up-regulated apoptosis-related pathways in the reperfusion stage. The main down-regulated pathways in endothelial cell clusters included *Blood vessel development* and other pathways in the reperfusion stage (Figure S5, Supporting Information). This phenomenon was similarly found in the overall stage with the reperfusion stage (Figure S6, Supporting Information). GSVA results indicated the dynamic changes of specific inflammatory pathways at different stages. In general, all endothelial clusters showed varying degrees of inflammatory activation after reperfusion. Pathways such as *NFKAPPAB_CANONICAL_PATHWAY, IL2_PI3K_PATHWAY, IL12_PATHWAY* and *IL23_PATHWAY* were significantly activated after reperfusion in almost all LSEC and VEC clusters (Figure 4D).

In order to investigate the potential intercellular communication network between mononuclear phagocyte and endothelial cell clusters after reperfusion, we analyzed the ligand-receptor pairs between these clusters (Figure 4E). Mononuclear phagocyte clusters commonly had *CD74_MIF, CD74_APP, C5AR1_RPS19* and *CCR1_CCL14* interactions with different endothelial cell clusters. The interactions between mononuclear phagocytes and LSEC clusters were highly enriched in *GRN_CLEC4M, ICAM3_CLEC4M, CCR1_CCL23* and other inflammatory cytokines ^24^ and cell adhesion related functions ^25^. Meanwhile, the interactions between mononuclear phagocytes and VEC clusters were enriched in nicotinamide phosphoribosyltransferase (NAMPT) related, *CD44_SELE* and other immune regulation related functions ^26,27^. Compared to TMo clusters, the interactions between KC and endothelial cell clusters were specifically enriched in *HLA–C_FAM3C, CCL4_SLC7A1* and *CD74_COPA*, most of which play an important role in IRI ^28-30^.

### 2.5 NK/T cells in graft IRI of liver transplantation

A total of 5844 single-cell transcriptomes were obtained from NK and T cell lineages of three samples and regrouped into 11 clusters (**Figure 5A**, Figure S7A, Supporting Information). Among them, there were four clusters of NK cells, which expressed high levels of *FCGR3A* and low levels of *CD3D*. The top five highly expressed genes were *GNLY, FGFBP2, FCGR3A, GZMB*, and *TYROBP* in GNLY_NK cluster; *PTGDS, FCER1G, MYOM2, AREG*, and *SPON2* in PTGDS_NK cluster; *XCL2, CMC1, KLRC3, KLRF1*, and *TYROBP* in XCL2_NK cluster; *XCL1, FCER1G, XCL2, AREG*, and *IL2RB* in XCL1_NK cluster. There were five clusters of T cell, including three CD8 T cell clusters, a CD4 T cell cluster and a γδ T cell cluster according to previous studies ^31,32^. We annotated them according to their characteristic expression profiles as *GZMK+* effect memory T cells (Tem), *FGFBP2+ NKG7+* effect T cells (Teff), and *SLC4A10+* mucosal-associated invariant T cells (MAIT). The top five highly expressed genes were *CD8B, GZMK, RGS1, CD8A*, and *COTL1* in CD8B_CD8 Tem; *CCL20, KLRB1, TRAV1-2, IL7R*, and *SLC4A10* in CCL20_CD8 MAIT; *IL7R, GPR183, LTB, RGCC*, and *LEF1* in IL7R_CD4 T; *TRBV9, TRAV38-2DV8, TRBV13, RP11-291B21*.*2*, and *TRGV5* in TRBV9_CD8 Teff; *TRDV2, TRGV9, KLRC1, TRDC* and *TRAC* in TRDV2_γδ T. For the other two clusters that highly expressed proliferation markers such as *MKI67* in comparison with other clusters, we annotated them as cycling (*CD8*+) T and (*FCGR3A+ CD3D-*) NK cells based on previous studies ^31,32^ (Figure 5A-B, Figure S7B, Table S12, Supporting Information).

**Figure 5:**
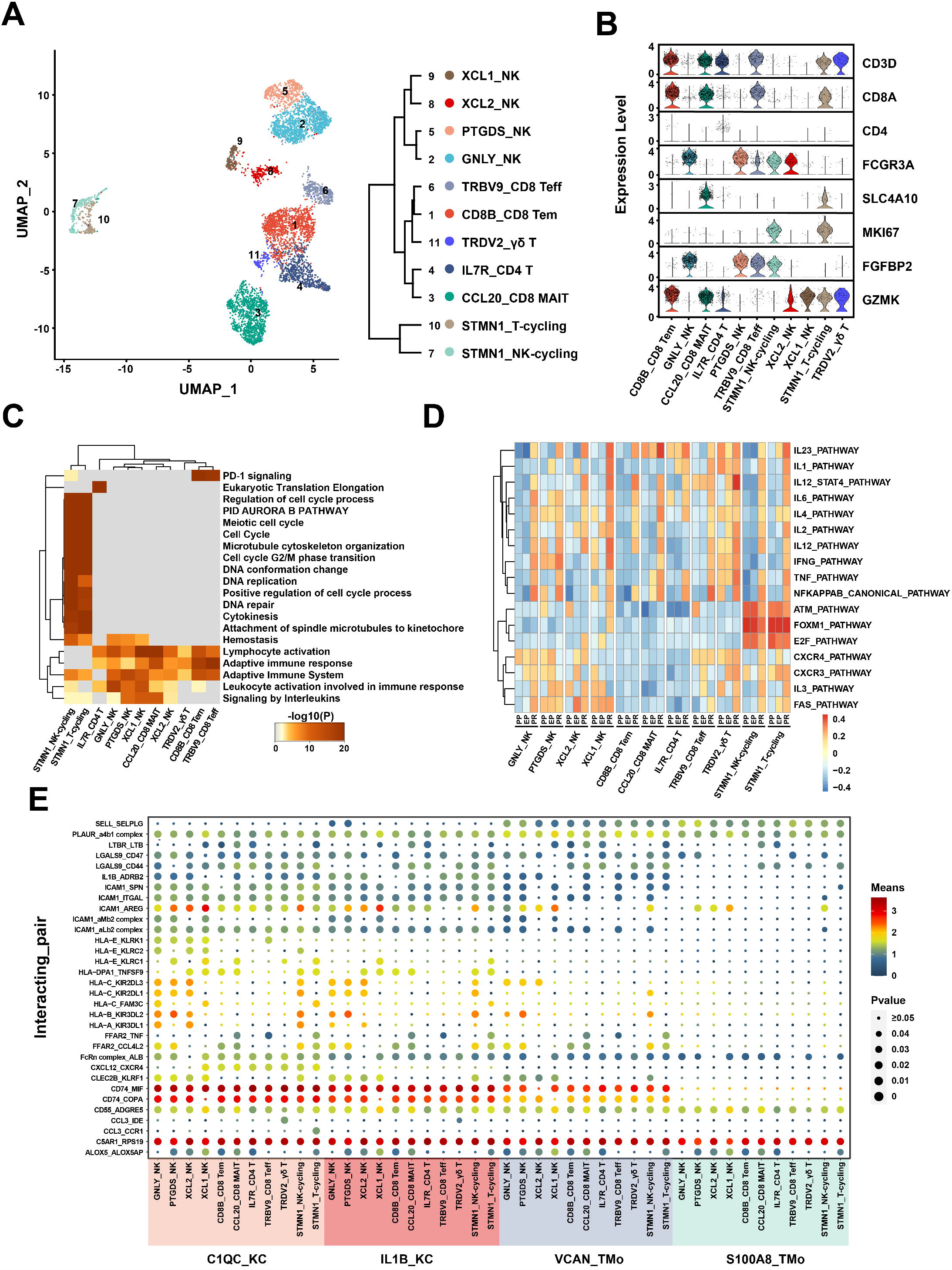
scRNA-seq of NK/T cells in liver transplantation. **A**: UMAP plot showing 11 NK/T cell clusters in liver transplantation, colored according to different clusters (left panel). Dendrogram of 11 clusters by the hierarchical clustering analysis based on their normalized mean expression values (right panel). **B**: Violin plots showing the normalized expression of marker genes (y axis) for NK/T cell clusters (x axis). **C**: Gene Ontology enrichment analysis results of NK/T cell clusters. Only the top 20 most significant GO terms (*P*-value < 0.05) are shown in rows. **D**: GSVA showing the pathways (PID gene sets) with significantly different activation in different samples of NK/T cell clusters. Different colors represent different activation scores. **E**: Cell-cell interaction analysis between mononuclear phagocyte clusters and different NK/T cell clusters in PR samples. Ligand-receptor pairs labeled in y-axis. The size of the circle represents the level of p-value, and different colors represent different means value. Ligands come from mononuclear phagocyte clusters, and receptors come from NK/T cell clusters.

The results of GO analysis showed that T and NK cell clusters are commonly enriched in the *Lymphocyte activation* and *Adaptive immune response* pathways, and the DEGs of NK cell clusters were mainly enriched in the *Phagocytosis* and *Regulation of reactive oxygen species metabolic process*. In T cell clusters, different clusters had different patterns. CD4 T cell cluster was specifically enriched in *Eukaryotic Translation Elongation*. CD8B_CD8 Tem and TRBV9_CD8 Teff cell clusters were specifically enriched in *PD-1 signaling*, while γδ T cell cluster rarely had a common enrichment pathway with other clusters. STMN1_T-cycling and STMN1_NK-cycling were clusters with high proliferation ability, and the DEGs were mainly enriched in *DNA replication* and *Positive regulation of cell cycle process* (Figure 5C, Figure S7C, Supporting Information).

In the study of the dynamic changes of NK and T cell clusters during liver transplantation, we found that IL7R_CD4 T, PTGDS_NK cells, TRBV9_CD8 Teff and CD8B_CD8 Tem clusters were more widely distributed in the PR sample than EP and PP samples (Figure S7D, Supporting Information). Compared with other clusters, CCL20_CD8 MAIT cluster expressed more DEGs in the reperfusion stage and GNLY_NK cluster expressed more DEGs in the cold preservation stage, indicating that CCL20_CD8 MAIT cluster was more sensitive to reperfusion injury and GNLY_NK cluster was more sensitive to cold storage injury. TRDV2_γδ T cluster had almost no DEGs during IRI, suggesting that TRDV2_γδ T cluster was not sensitive to IRI (Figure S7E, Supporting Information). The amount of up-regulated DEGs in overall stage was slightly more than that in reperfusion stage, and they shared most of the genes especially with high expression levels (Figure S7F, Table S13-15, Supporting Information).

The GO analysis of DEGs in EP and PR samples showed that most of cell clusters were activated in *TNF signaling pathway, NF-kappa B signaling pathway, Myeloid leukocyte activation, T cell activation* and other pathways related to inflammation and cell activation in the reperfusion stage (Figure S8A-B). GSVA was performed to observe the dynamic changes of specific inflammatory pathways. We found that inflammation-related pathways such as *IL23_PATHWAY, IL6_PATHWAY, TNF_PATHWAY* and *NFKAPPAB_CANONICAL_PATHWAY* were obviously activated after reperfusion in most NK and T cell clusters. Cell cycle-related pathways such as *ATM_PATHWAY, FOXM1_PATHWAY* and *E2F_PATHWAY* were mainly expressed in cycling cell clusters. Pathways such as *FAS_PATHWAY, CXCR4_PATHWAY* and *CXCR3_PATHWAY* were mainly expressed in most NK and T cell clusters except for CD8B_CD8 Tem, CCL20_CD8 MAIT and IL7R_CD4 T cell clusters (Figure 5D).

Mononuclear phagocyte clusters had universal interactions with NK/T cell clusters in cell-cell interaction analysis (Figure 5E), including C5AR1_RPS19, CD55_ADGRE5, PLAUR_a4b1 complex and other ligand-receptor pairs. Compared with S100A8_TMo cluster, VCAN_TMo cluster was enriched in ligand-receptor pairs such as CD74_MIF, CD74_COPA and ICAM1_ITGAL in the interaction with NK and T cell clusters. The interactions between NK cell and KC cell clusters were specifically enriched in human leukocyte antigen (HLA) related and killer cell immunoglobulin-like receptors (KIR) related receptor ligand pairs such as HLA–C_KIR2DL3 and HLA–B_KIR3DL2. The interactions between T cell and KC cell clusters (especially in C1QC_KC) were mainly enriched in chemokine ligand related receptor ligand pairs such as CXCL12_CXCR4, indicating the role of immune surveillance and T cell recruitment of C1QC_KC cluster ^33^.

### 2.6 B and plasma cells in graft IRI of liver transplantation

We obtained the single cell transcriptomes of 1,796 cells from B and plasma cells lineage of three samples and re-clustered them into six clusters (**Figure 6A**, Figure S9A, Supporting Information). These clusters uncovered three *MS4A1+* B cell clusters and three *SDC1+* plasma cell clusters in the liver with different canonical marker genes. The top five highly expressed genes in the B cell clusters were *TCL1A, TXNIP, BTG1, CD37*, and *FCER2* in TCL1A_B cell cluster; *GPR183, TNF, COTL1, AC079767*, and *NR4A2* in GPR183_B cell cluster; *MT2A, FGR, FCRL3, FCRL5*, and *CIB1* in MT2A_B cell cluster. The top five highly expressed genes in the three plasma clusters were *IGKV1D-16, IGKV3D-20, IGKV1-16, IGHV3-72*, and *IGKV2D-28* in IGKV1D-16_plasma cell cluster; *HIST1H4C, RRM2, GAPDH, HMGB2*, and *IGHV6-1* in HIST1H4C_plasma cell cluster; *IGLV3-1, IGKV1D-39, CH17-224D4*.*2, IGKV1D-12*, and *IGLV3-21* in IGLV3-1_plasma cell cluster (Figure 6B, Figure S9B, Table S16, Supporting Information).

**Figure 6:**
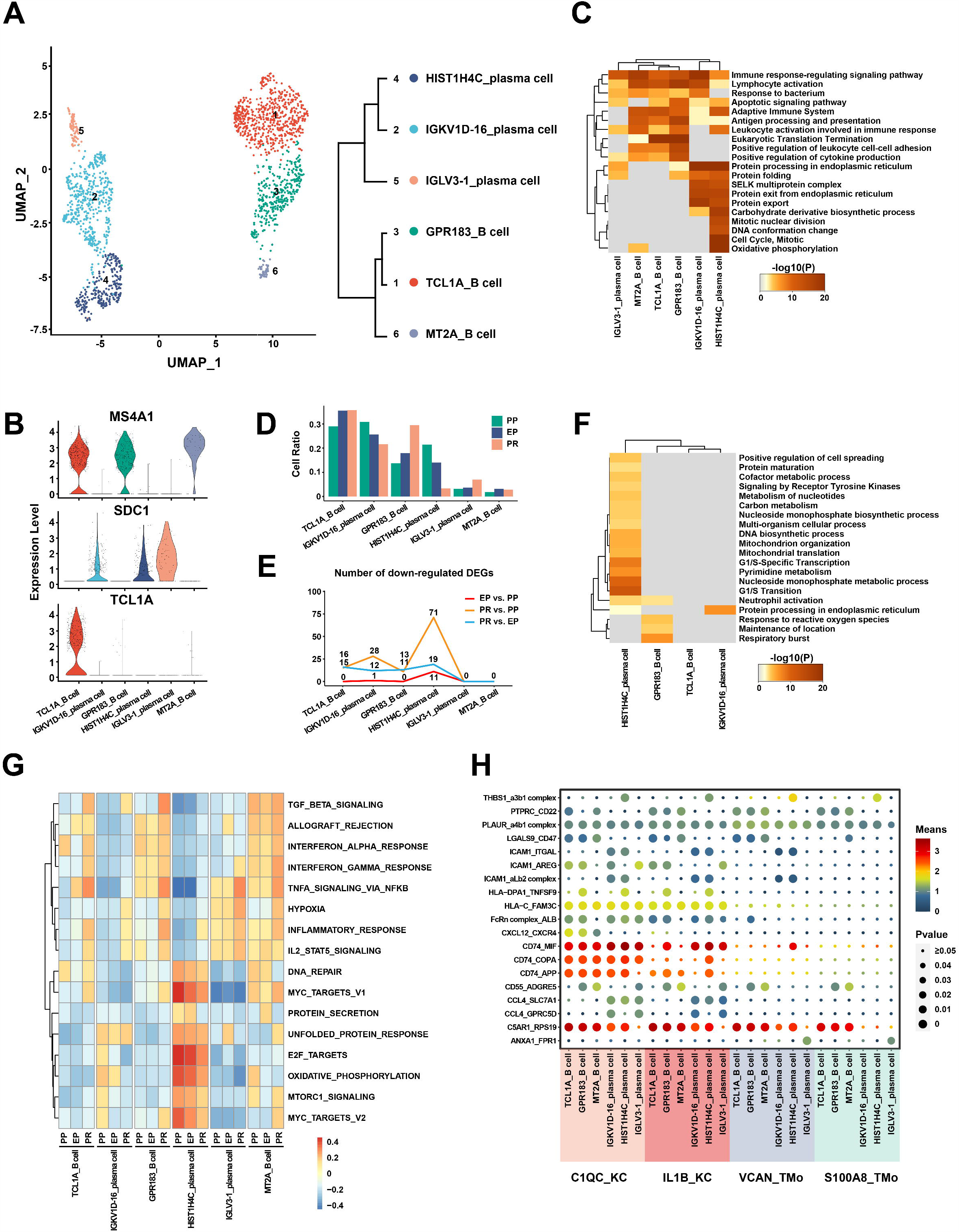
scRNA-seq of B/plasma cells in liver transplantation. **A**: UMAP plot showing six B/plasma cell clusters in liver transplantation, colored according to different clusters (left panel). Dendrogram of six clusters by hierarchical clustering analysis based on their normalized mean expression values (right panel). **B**: Violin plots showing the normalized expression of *MS4A1, SDC1* and *TCL1A* genes (y axis) for B/plasma cell clusters (x axis). **C**: Gene Ontology enrichment analysis results of B/plasma cell clusters. Only the top 20 most significant GO terms (*P*-value < 0.05) are shown in rows. **D**: Cell ratio of different B/plasma cell clusters in PP, EP and PR samples. GSVA showing the pathways (PID gene sets) with significantly different activation in different samples of B/plasma cell clusters. Different colors represent different activation scores. **E**: Number of down-regulated DEGs between different samples in different B/plasma cell clusters. **F**: Gene Ontology enrichment analysis results of down-regulated DEGs from PR vs. PP. Only the top 20 most significant GO terms (P-value < 0.05) are shown in rows. **G**: GSVA showing the pathways (HALLMARK gene sets) with significantly different activation in different samples of B/plasma cell clusters. Different colors represent different activation scores. **H**: Cell-cell interaction analysis between mononuclear phagocyte clusters and different B/plasma cell clusters in PR samples. Ligand-receptor pairs labeled in y-axis. The size of the circle represents the level of p-value, and different colors represent different means value. Ligands come from mononuclear phagocyte clusters, and receptors come from B/plasma cell clusters.

The GO analysis results showed that different B and plasma cell clusters have different physiological functions. The common enrichment pathways of B cell and plasma cell clusters included *Lymphocyte activation, Immune response-regulating signaling pathway, Adaptive Immune System* and so on. In addition, the B cell clusters were specifically enriched in the *Eukaryotic Translation Termination* and *Positive regulation of leukocyte cell-cell adhesion* pathways while the plasma cell clusters were mainly enriched in the *Protein processing in endoplasmic reticulum* and *Protein export* pathways. We observed that *HIST1H4C*_plasma cell cluster was also specifically enriched in *Cell Cycle, DNA conformation change* and *mitotic nuclear division* pathways, indicating that this cluster may have active proliferation function (Figure 6C).

In the study of dynamic changes at different stages of liver transplantation, we found that the number of DEGs in B and plasma cell clusters during the cold preservation stage and reperfusion stage was less than that of other cell lineages, suggesting that the expression profile of B and plasma cell clusters was not obviously altered after IRI (Figure S9C). Interestingly, we found that the number of down-regulated genes in the *HIST1H4C*_plasma cell cluster increased during the cold preservation and reperfusion stage (Figure 6E). These down-regulated genes were mainly enriched in cell cycle-related pathways such as *G1/S Transition, G1/S-Specific Transcription* and *mitochondrial translation* (Figure 6F, Figure S9D-F, Table S17-19). At the same time, the results of GSVA analysis showed that the expression of genes in HIST1H4C_plasma cell cluster related to *E2F_TARGETS, G2M_CHECKPOINT* and *PROTEIN_SECRETION* pathways gradually decreased during the cold preservation and reperfusion stage (Figure 6G). Combined with the proportional decrease in the cell number of HIST1H4C_plasma cell cluster throughout the PP, EP and PR samples, we speculate that IRI can affect the cell cycle and reduce the cell proliferation of HIST1H4C_plasma cell cluster (Figure 6D). In addition, we found that the APS of plasma cells was lower than that of B cells in MHC class II genes (*P*<0.0001), and the APS of HIST1H4C_plasma cell cluster decreased significantly after reperfusion (*P*=0.0002).

In the study of the cell-cell interaction between mononuclear phagocyte and B plasma cell clusters after reperfusion, we found that the ligand-receptor pairs such as *PLAUR_a4b1 complex* and *C5AR1_RPS19* were commonly enriched between mononuclear phagocyte and B plasma cell clusters. The interactions between mononuclear phagocytes and B cells were specifically enriched in *LGALS9_CD47* and *PTPRC_CD22* and other cell-cell/cell-matrix interactions related functions ^34,35^. In addition, ligand-receptor pairs such as *HLA–C_FAM3C, CD74_MIF, CD74_COPA, CD74_APP* which play an important role in IRI, were enriched in the interaction between KC and B plasma cell clusters ^28-30^. Compare to the interactions between KC and B cell clusters, those between KC and plasma cell clusters were specifically enriched in *CCL4_SLC7A1* and *CCL4_GPRC5D* which may relate to possible chemotaxis function in KC clusters ^36,37^ (Figure 6H).

## 3 DISCUSSION

To the best of our knowledge, this study obtained the first unbiased and comprehensive liver transplant cell atlas by using scRNA-seq. We annotated the transcriptome characteristics of intrahepatic mononuclear phagocyte, endothelial, NK, T, B and plasma cell clusters in details and revealed the dynamic changes of transcriptome in each cluster during IRI. In addition, we identified the potential interactions between mononuclear phagocyte clusters and other cell clusters after graft reperfusion.

Excessive inflammation caused by KCs is the key mechanism that causes pathological damage in IRI. After reperfusion, the accumulated endogenous damage associated molecular patterns (DAMPs) and pathogen associated molecular patterns (PAMPs), are released into the liver, which activate KCs, and produce ROS, TNF-α, IL-1β and other pro-inflammatory cytokines, forming a positive feedback loop ^8,38^. At the same time, these cytokines lead to the expression of adhesion molecules on the surface of LSEC and recruitment of monocytes, neutrophils and T cells ^39,40^. On the other hand, some studies have reported that KCs can reduce liver IRI damage by producing nitric oxide (NO) to relax blood vessels, up-regulate the expression of hemeoxygenase 1 to eliminate pro-inflammatory factors and produce IL-10 ^17,41-43^. MacParland et al had observed the existence of macrophages with different functions in fresh liver tissue, including the *CD68+MARCO+* cell subpopulation annotated as KCs with inflammation-regulating properties, and the *CD68+MARCO-*cell subpopulation annotated as a recently recruited macrophages with pro-inflammatory properties ^14^. However, how KC implements these different functions has not yet been well explained. The *CD68+MARCO+/-*cell types were also found in our liver transplantation scRNA-seq dataset (Figure S1D). Our study clearly defined two populations of KC clusters with different functions through scRNA-seq, which might explain the different pro-inflammatory and anti-inflammatory functions of KC in IRI. In this study, we demonstrated a possible protective effect of *TNIP3* expression in KC clusters during IRI. In recent studies, Liu D etc. confirmed that adenovirus-mediated TNIP3 expression in the liver blocked NASH progression in mice, indicating that TNIP3 may be a promising therapeutic target for NASH ^44^. TNIP3 can bind to zinc finger protein TNFAIP3 and inhibit NF-κB activation induced by TNF, Toll-like receptor 4 (TLR4), interleukin-1 and 12-O-tetradecanoyl phorbol-13-acetate ^45^. Consistent with previous studies ^46,47^, the NF-κB pathway was generally activated after reperfusion in mononuclear phagocyte clusters in our dataset. At the same time, we observed that KC clusters specifically showed increased expression of *TNIP3* after reperfusion and the *TNIP3* high expression group demonstrated reduced level of postoperative liver injury. These results indicated that the inhibition of NF-κB pathway activation by TNIP3 may be one of the mechanisms by which KC cluster plays a protective role against graft IRI during liver transplantation.

We discovered the potential interaction pathways between mononuclear phagocytes and four LSEC clusters in our study. In the current dataset, four LSEC clusters specifically expressed *CLEC4M* and might receive *GRN* signaling molecules from mononuclear phagocytes. In previous studies, promoting *GRN* expression can reduce brain IRI-induced brain damage by inhibiting neuronal apoptosis and ROS production ^48^; recombinant-GRN treatment can inhibit the recruitment of neutrophils and reduced the activation of NF-κB and MMP-9 in the brain IRI, and GRN has been reported to have a protective effect on inflammation caused by kidney and brain IRI ^49,50^. Therefore, mononuclear phagocytes may express *GRN* and send a signal to the *CLEC4M* receptor on LSEC to reduce the recruitment of neutrophils, thereby reducing liver IRI. In addition, we discovered a special interaction mechanism between the C1QC_KC cluster and specific T and B cell clusters. Previous studies have confirmed that the expression of *CXCL12* increases in damaged liver tissue and contributes to recruitment of *CXCR4+* cells ^51^. In our study, compared to other mononuclear phagocyte clusters, the C1QC_KC cluster specifically expresses *CXCL12* and sends a signal to the *CXCR4* receptor expressed on the T cell and B cell clusters, and increases the migration ability of the corresponding cell clusters.

We acknowledge that the current research has some limitations. Firstly, we did not obtained a sufficient number of hepatocytes and bile duct epithelial cells, which also play a very important role in IRI. The hepatocyte populations are particularly susceptible to dissociation. Nowadays, the collagenase perfusion method is routinely used to separate hepatocytes, but due to the need to obtain liver tissues from the same donor at different time points and the limited sample size, we cannot use the collagenase perfusion method to separate cells. Secondly, our samples have no biological replicates, even though we have obtained a sufficient number of cells, a larger sample size will strengthen our conclusions. In terms of important molecular mechanisms, we combined bulk RNA-seq data and immunofluorescence or Western Blot experiments to verify our conclusions.

Overall, we have obtained the first unbiased and comprehensive liver transplant cell atlas. We annotated the subpopulations of multiple cell types and described the dynamic changes of their transcriptome in IRI and the interaction of mononuclear phagocyte clusters with other cell clusters after reperfusion, which allows us to investigate the mechanism of IRI from single-cell resolution. At the same time, the *TNIP3* gene which is specifically up-regulated in KC clusters after reperfusion, may be a potential therapeutic target of IRI.

## 4 MATERIALS AND METHODS

### 4.1 Human liver tissue collection and dissociation

All PP, EP, PR liver tissues were obtained from a 47-year-old male brain death donor. The recipient was a 51-year-old man with chronic hepatitis B virus infection and hepatocellular carcinoma. This study protocol was approved by the Ethics Committee of the First Affiliated Hospital of Sun Yat-sen University (permit no. [2018]255) and comply with the Declaration of Helsinki principles.

Blood or cold preservation solution (University of Wisconsin (UW) solution) in the liver tissues were immediately washed away with 4°C physiological saline after tissue collection. The tissues were cut into 3 mm pieces, incubated with 1 mM EGTA Sigma-Aldrich, Cat. no. E0396-10G) and rotate at 37°C, 50 rpm for 10 minutes. After washing away the EGTA, the tissues were treated in the digestion solution (1 mg/ml Collagenase II + 1 mg/ml Collagenase IV + 50 ug/ml DNase I) at 100 rpm and 37 °C for 30 minutes. The cell suspension was passed through a 70 mm nylon cell strainer (BD, Cat. No. 352350), and then centrifuged at 50×g for 3 minutes to collect the cell pellet, and then the remaining suspension centrifuged at 300×g for 5 minutes to collect the remaining cell pellet. Each tissue was resuspended to a concentration of 50-500 million cells per milliliter with resuspension buffer. Then used LIVE/DEAD Viability/Cytotoxicity Kit (Invitrogen, Cat. no. L3224) to stained cells, and only AM+EH-cells were collected by FACS for each tissue.

### 4.2 Single cell RNA-seq data processing

Single-cell cDNA libraries were constructed using the standard procedure of GemCode technology (10X Genomics, USA). For each tissue, the 5’-end cDNA libraries were sequenced using data volumes of 8,000 cells. Illumina HiSeq XTen platform was used for sequencing at pared-end 150 bp length. The cDNA library measured 120G base data volume.

The Illumina software bcl2fastq (version v2.19.0.316) was used to convert the raw data (BCL files) into fastq files. CellRanger (version 3.0.1) was used to count to align the sequence to the human reference genome (hg38) and calculate the gene expression matrix. The original gene expression matrix in the “filtered_feature_bc_matrix” folder generated by CellRanger software was used for further analysis.

### 4.3 Doublets classifying, cell clustering, and differential gene expression analysis

Data filtering, normalizing, dimensionality reduction and clustering for cells were performed using R software (version 3.6.1; https://www.r-project.org) and Seurat package (version 3.1.2; https://satijalab.org/seurat). “DoubletFinder” (version 2.0.3; https://github.com/chris-mcginnis-ucsf/DoubletFinder) was used to identify doublets in each sample. A further method of cell filtration is to remove cells with low quality (UMI < 1,000, gene number < 500), and high (> 0.25) mitochondrial genome transcript ratio. In the analysis of each cell type, a subpopulation of cells expressing multiple markers of different cell types will be defined as a non-single cell population and will be removed in the subsequent analysis. “NormalizedData” function was used to normalize the expression data, and then used the “ScaleData” function to perform regression to remove the influence of UMI and mitochondrial content. “FindVariableGenes” was used to identify genes with high variation for principal component analysis (PCA analysis). For each sample and main cell type, we used a different number of principal components (PC) and resolution for dimensionality reduction analysis. Use the same number of PCs to identify cell types as in UMAP dimensionality reduction (“FindClusters” function). “FindAllMarkers” function was used to analyze the differential expression markers, and use Wilcoxon to test the significance level. Genes with an absolute value of FC (fold-change) natural number logarithm (|lnFC|) greater than 0.25 and a *P*-adjust value lower than 0.05 after adjustment by Bonferroni are considered to have significant differences.

### 4.4 Analysis of pathway and cellular interaction analysis

Gene Ontology (GO) enrichment analysis was performed using the online tools Metascape (http://metascape.org/gp/index.html) ^52^. Genes with LnFC greater than 0.405 and *P*-adjust value less than 0.05 were selected for GO enrichment analysis. Gene set variation analysis (GSVA) was performed to identify enriched cellular pathways in our dataset, using R package GSVA (version 1.32.0) (https://www.gsea-msigdb.org/gsea/index.jsp) on the C2: Canonical pathways-BIOCARTA subset (c2.cp.biocarta.v7.1.symbols.gmt), PID subset (c2.cp.pid.v7.1.symbols.gmt) and 50 hallmark pathways with default parameters. “AddModuleScore” function in the R Seurat package was used to calculate the antigen presentation scores related to MHC I and MHC II molecules according to REACTOME database geneset.

### 4.5 Immunofluorescence and Western Blot

The paraffin-embedded tissue sections were deparaffinized, rehydrated and treated with 3% H2O2 to block endogenous peroxidase activity and underwent high-temperature antigen retrieval. The repaired tissue was incubated with 3% BSA for 30 minutes at room temperature, and incubated overnight with CD68 antibody (Servicebio, GB11067) in a refrigerator at 4 °C. Then the slides were incubated with the secondary antibody (HRP polymer, anti-rabbit IgG) for 50 minutes at room temperature. Subsequently, the tissue was treated with a solution containing TSA reagent for 10 minutes. After each treatment with TSA, the slides were subjected to microwave heat treatment, incubated overnight with TNIP3 antibody (Abcam, ab198697) in a refrigerator at 4 °C, and then incubated with the secondary antibody and treated with TSA. For each sample, after all antigen labeling was completed, the nucleus was stained with 4’-6’-di-yl-2-phenylindole (DAPI).

Total protein was extracted using RIPR lysis buffer (KeyGEN, China) and quantified using the BCA protein assay (Thermo Scientific). The proteins of tissue samples were resolved by 10% SDS-PAGE and transferred to the PVDF membrane (0.2 µm pore size, Millipore, Billerica, MA, USA). The membranes were immunoblotted with primary antibodies overnight at 4°C followed by secondary antibody incubation for 2 hours, and subsequent visualization with Chemiluminescent HRP substrate (Millipore). The antibodies were used as follows: TNIP3 (1:2,000, Abcam, ab198697); β-tubulin (1:2,000, cat. No. 100941-1-AP; Proteintech); Goat anti-Rabbit IgG antibody (1:5,000, cat. No. ab205718; Abcam).

## Data sharing and code availability

The key raw data have been deposited in the Gene Expression Omnibus (GEO; https://www.ncbi.nlm.nih.gov/geo/). The rest of the data and code are available from the authors upon reasonable request.

## Conflict of Interests

The authors declare no conflict of interest.

## AUTHORSHIP CONTRIBUTIONS

X.S.H., Z.Y.G. and J.X.B. designed the study; L.H.W., S.H., H.T.C, D.W.Z. and J.H.X. did the sample collection, library construction, and data generation; L.H.W. analyzed the scRNA-seq data; L.H.W, J.L., S.R.C., T.L., X.Y.Y. and S.J.H. did the immunofluorescence and Western Blot experiments; L.H.W., Z.Y.G. and X.S.H. wrote the paper.

## ACKNOWLEDGEMENTS

This work was supported by grants from the National Natural Science Foundation of China (81970564 to XSH, 81471583 to ZYG and 81570587 to ZYG), Guangdong Provincial Key Laboratory of Organ Donation and Transplant Immunology, The First Affiliated Hospital, Sun Yat-sen University, Guangzhou, China (2013A061401007 to XSH and 2017B030314018 to XSH), Guangdong Provincial International Cooperation Base of Science and Technology (Organ Transplantation), The First Affiliated Hospital, Sun Yat-sen University, Guangzhou, China (2015B050501002 to XSH), Guangdong Provincial Natural Science Funds for Major Basic Science Culture Project (2015A030308010 to XSH), Special Support Program for Training High Level Talents in Guangdong Province (2015TQ01R168 to ZYG), Pearl River Nova Program of Guangzhou (201506010014 to ZYG).

## Notes

### Competing Interest Statement

The authors have declared no competing interest.

