## Supplementary Figures for "Resolving the graft ischemia-reperfusion injury during liver transplantation at the single cell resolution"

### Figure S1

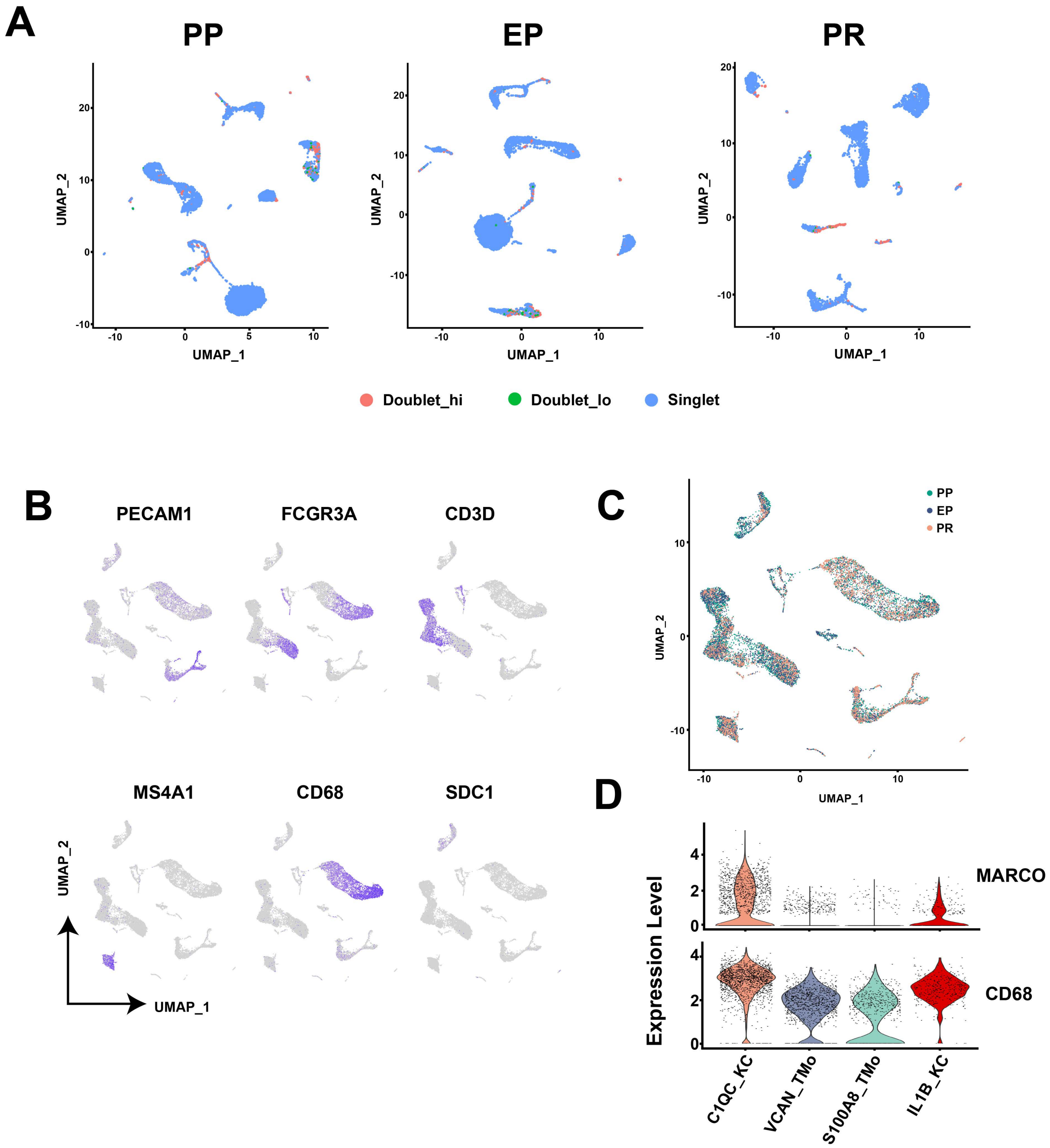

A
PP
EP
PR
UMAP_1
UMAP_2
Doublet_hi
Doublet_lo
Singlet
B
PECAM1
FCGR3A
CD3D
MS4A1
CD68
SDC1
UMAP_2
UMAP_1
C
PP
EP
PR
UMAP_2
UMAP_1
D
Expression Level
MARCO
CD68
C1QC_KC
VCAN_TMo
S100A8_TMo
IL1B_KC

### Figure S2

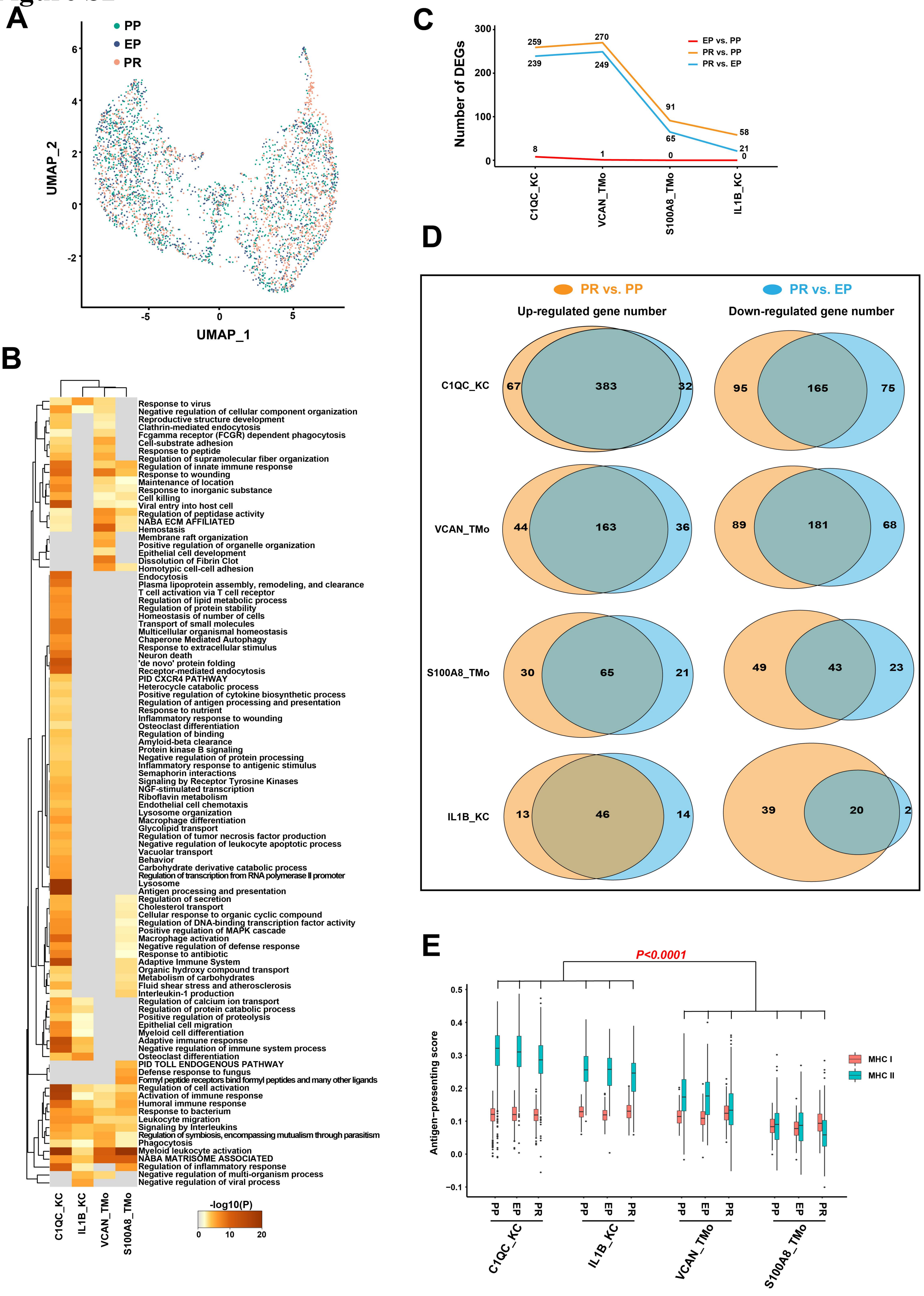

A
PP
EP
PR
UMAP_2
UMAP_1
B
-log10(P)
Response to virus
Negative regulation of cellular component organization
Reproductive structure development
Clathrin-mediated endocytosis
Fcgamma receptor (FCGR) dependent phagocytosis
Cell-substrate adhesion
Response to peptide
Regulation of supramolecular fiber organization
Regulation of innate immune response
Response to wounding
Maintenance of location
Response to inorganic substance
Cell killing
Viral entry into host cell
Regulation of peptidase activity
NABA ECM AFFILIATED
Hemostasis
Membrane raft organization
Positive regulation of organelle organization
Epithelial cell development
Dissolution of Fibrin Clot
Homotypic cell-cell adhesion
Endocytosis
Plasma lipoprotein assembly, remodeling, and clearance
T cell activation via T cell receptor
Regulation of lipid metabolic process
Regulation of protein stability
Homeostasis of number of cells
Transport of small molecules
Multicellular organismal homeostasis
Chaperone Mediated Autophagy
Response to extracellular stimulus
Neuron death
'de novo' protein folding
Receptor-mediated endocytosis
PID CXCR4 PATHWAY
Heterocycle catabolic process
Positive regulation of cytokine biosynthetic process
Regulation of antigen processing and presentation
Response to nutrient
Inflammatory response to wounding
Osteoclast differentiation
Regulation of binding
Amyloid-beta clearance
Protein kinase B signaling
Negative regulation of protein processing
Inflammatory response to antigenic stimulus
Semaphorin interactions
Signaling by Receptor Tyrosine Kinases
NGF-stimulated transcription
Riboflavin metabolism
Endothelial cell chemotaxis
Lysosome organization
Macrophage differentiation
Glycolipid transport
Regulation of tumor necrosis factor production
Negative regulation of leukocyte apoptotic process
Vacuolar transport
Behavior
Carbohydrate derivative catabolic process
Regulation of transcription from RNA polymerase II promoter
Lysosome
Antigen processing and presentation
Regulation of secretion
Cholesterol transport
Cellular response to organic cyclic compound
Regulation of DNA-binding transcription factor activity
Positive regulation of MAPK cascade
Macrophage activation
Negative regulation of defense response
Response to antibiotic
Adaptive Immune System
Organic hydroxy compound transport
Metabolism of carbohydrates
Fluid shear stress and atherosclerosis
Interleukin-1 production
Regulation of calcium ion transport
Regulation of protein catabolic process
Positive regulation of proteolysis
Epithelial cell migration
Myeloid cell differentiation
Adaptive immune response
Negative regulation of immune system process
Osteoclast differentiation
PID TOLL ENDOGENOUS PATHWAY
Defense response to fungus
Formyl peptide receptors bind formyl peptides and many other ligands
Regulation of cell activation
Activation of immune response
Humoral immune response
Response to bacterium
Leukocyte migration
Signaling by Interleukins
Regulation of symbiosis, encompassing mutualism through parasitism
Phagocytosis
Myeloid leukocyte activation
NABA MATRISOME ASSOCIATED
Regulation of inflammatory response
Negative regulation of multi-organism process
Negative regulation of viral process
C1QC_KC
IL1B_KC
VCAN_TMo
S100A8_TMo
C
Number of DEGs
EP vs. PP
PR vs. PP
PR vs. EP
259
270
239
249
91
65
58
21
8
1
0
0
C1QC_KC
VCAN_TMo
S100A8_TMo
IL1B_KC
D
PR vs. PP
PR vs. EP
Up-regulated gene number
Down-regulated gene number
C1QC_KC
67
383
32
95
165
75
VCAN_TMo
44
163
36
89
181
68
S100A8_TMo
30
65
21
49
43
23
IL1B_KC
13
46
14
39
20
2
E
P<0.0001
Antigen−presenting score
MHC I
MHC II
PP
EP
PR
C1QC_KC
IL1B_KC
VCAN_TMo
S100A8_TMo

Figure S3

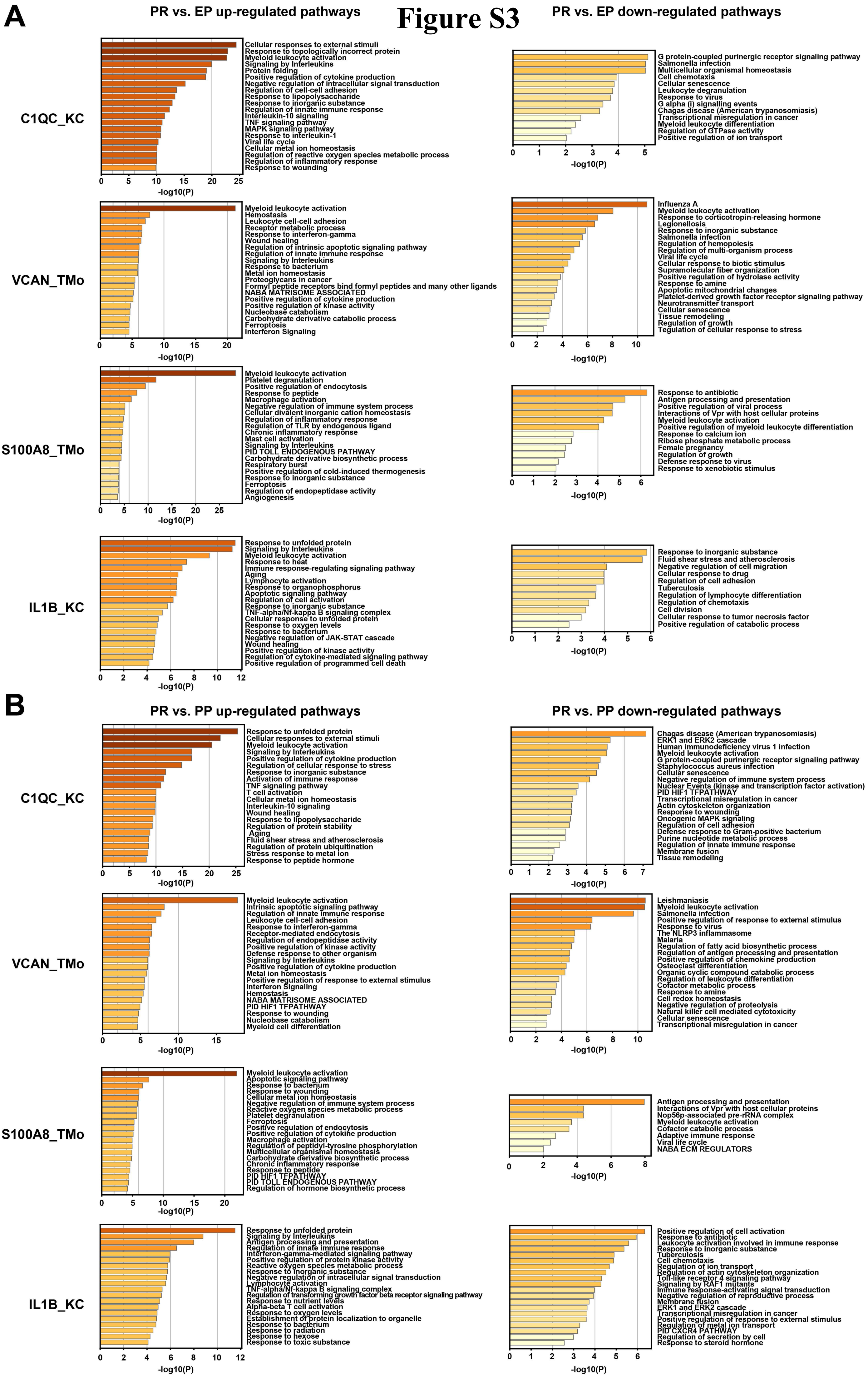
A
PR vs. EP up-regulated pathways
PR vs. EP down-regulated pathways
C1QC_KC
Cellular responses to external stimuli
Response to topologically incorrect protein
Myeloid leukocyte activation
Signaling by Interleukins
Protein folding
Positive regulation of cytokine production
Negative regulation of intracellular signal transduction
Regulation of cell-cell adhesion
Response to lipopolysaccharide
Response to inorganic substance
Regulation of innate immune response
Interleukin-10 signaling
TNF signaling pathway
MAPK signaling pathway
Response to interleukin-1
Viral life cycle
Cellular metal ion homeostasis
Regulation of reactive oxygen species metabolic process
Regulation of inflammatory response
Response to wounding
-log10(P)
G protein-coupled purinergic receptor signaling pathway
Salmonella infection
Multicellular organismal homeostasis
Cell chemotaxis
Cellular senescence
Leukocyte degranulation
Response to virus
G alpha (i) signalling events
Chagas disease (American trypanosomiasis)
Transcriptional misregulation in cancer
Myeloid leukocyte differentiation
Regulation of GTPase activity
Positive regulation of ion transport
VCAN_TMo
Myeloid leukocyte activation
Hemostasis
Leukocyte cell-cell adhesion
Receptor metabolic process
Response to interferon-gamma
Wound healing
Regulation of intrinsic apoptotic signaling pathway
Regulation of innate immune response
Signaling by Interleukins
Response to bacterium
Metal ion homeostasis
Proteoglycans in cancer
Formyl peptide receptors bind formyl peptides and many other ligands
NABA MATRISOME ASSOCIATED
Positive regulation of cytokine production
Positive regulation of kinase activity
Nucleobase catabolism
Carbohydrate derivative catabolic process
Ferroptosis
Interferon Signaling
Influenza A
Myeloid leukocyte activation
Response to corticotropin-releasing hormone
Legionellosis
Response to inorganic substance
Salmonella infection
Regulation of hemopoiesis
Regulation of multi-organism process
Viral life cycle
Cellular response to biotic stimulus
Supramolecular fiber organization
Positive regulation of hydrolase activity
Response to amine
Apoptotic mitochondrial changes
Platelet-derived growth factor receptor signaling pathway
Neurotransmitter transport
Cellular senescence
Tissue remodeling
Regulation of growth
Tegulation of cellular response to stress
S100A8_TMo
Myeloid leukocyte activation
Platelet degranulation
Positive regulation of endocytosis
Response to peptide
Macrophage activation
Negative regulation of immune system process
Cellular divalent inorganic cation homeostasis
Regulation of inflammatory response
Regulation of TLR by endogenous ligand
Chronic inflammatory response
Mast cell activation
Signaling by Interleukins
PID TOLL ENDOGENOUS PATHWAY
Carbohydrate derivative biosynthetic process
Respiratory burst
Positive regulation of cold-induced thermogenesis
Response to inorganic substance
Ferroptosis
Regulation of endopeptidase activity
Angiogenesis
Response to antibiotic
Antigen processing and presentation
Positive regulation of viral process
Interactions of Vpr with host cellular proteins
Myeloid leukocyte activation
Positive regulation of myeloid leukocyte differentiation
Response to calcium ion
Ribose phosphate metabolic process
Female pregnancy
Regulation of growth
Defense response to virus
Response to xenobiotic stimulus
IL1B_KC
Response to unfolded protein
Signaling by Interleukins
Myeloid leukocyte activation
Response to heat
Immune response-regulating signaling pathway
Aging
Lymphocyte activation
Response to organophosphorus
Apoptotic signaling pathway
Regulation of cell activation
Response to inorganic substance
TNF-alpha/Nf-kappa B signaling complex
Cellular response to unfolded protein
Response to oxygen levels
Response to bacterium
Negative regulation of JAK-STAT cascade
Wound healing
Positive regulation of kinase activity
Regulation of cytokine-mediated signaling pathway
Positive regulation of programmed cell death
Response to inorganic substance
Fluid shear stress and atherosclerosis
Negative regulation of cell migration
Cellular response to drug
Regulation of cell adhesion
Tuberculosis
Regulation of lymphocyte differentiation
Regulation of chemotaxis
Cell division
Cellular response to tumor necrosis factor
Positive regulation of catabolic process
B
PR vs. PP up-regulated pathways
PR vs. PP down-regulated pathways
C1QC_KC
Response to unfolded protein
Cellular responses to external stimuli
Myeloid leukocyte activation
Signaling by Interleukins
Positive regulation of cytokine production
Regulation of cellular response to stress
Response to inorganic substance
Activation of immune response
TNF signaling pathway
T cell activation
Cellular metal ion homeostasis
Interleukin-10 signaling
Wound healing
Response to lipopolysaccharide
Regulation of protein stability
Aging
Fluid shear stress and atherosclerosis
Regulation of protein ubiquitination
Stress response to metal ion
Response to peptide hormone
Chagas disease (American trypanosomiasis)
ERK1 and ERK2 cascade
Human immunodeficiency virus 1 infection
Myeloid leukocyte activation
G protein-coupled purinergic receptor signaling pathway
Staphylococcus aureus infection
Cellular senescence
Negative regulation of immune system process
Nuclear Events (kinase and transcription factor activation)
PID HIF1 TFPATHWAY
Transcriptional misregulation in cancer
Actin cytoskeleton organization
Response to wounding
Oncogenic MAPK signaling
Regulation of cell adhesion
Defense response to Gram-positive bacterium
Purine nucleotide metabolic process
Regulation of innate immune response
Membrane fusion
Tissue remodeling
VCAN_TMo
Myeloid leukocyte activation
Intrinsic apoptotic signaling pathway
Regulation of innate immune response
Leukocyte cell-cell adhesion
Response to interferon-gamma
Receptor-mediated endocytosis
Regulation of endopeptidase activity
Positive regulation of kinase activity
Defense response to other organism
Signaling by Interleukins
Positive regulation of cytokine production
Metal ion homeostasis
Positive regulation of response to external stimulus
Interferon Signaling
Hemostasis
NABA MATRISOME ASSOCIATED
PID HIF1 TFPATHWAY
Response to wounding
Nucleobase catabolism
Myeloid cell differentiation
Leishmaniasis
Myeloid leukocyte activation
Salmonella infection
Positive regulation of response to external stimulus
Response to virus
The NLRP3 inflammasome
Malaria
Regulation of fatty acid biosynthetic process
Regulation of antigen processing and presentation
Positive regulation of chemokine production
Osteoclast differentiation
Organic cyclic compound catabolic process
Regulation of leukocyte differentiation
Cofactor metabolic process
Response to amine
Cell redox homeostasis
Negative regulation of proteolysis
Natural killer cell mediated cytotoxicity
Cellular senescence
Transcriptional misregulation in cancer
S100A8_TMo
Myeloid leukocyte activation
Apoptotic signaling pathway
Response to bacterium
Response to wounding
Cellular metal ion homeostasis
Negative regulation of immune system process
Reactive oxygen species metabolic process
Platelet degranulation
Ferroptosis
Positive regulation of endocytosis
Positive regulation of cytokine production
Macrophage activation
Regulation of peptidyl-tyrosine phosphorylation
Multicellular organismal homeostasis
Carbohydrate derivative biosynthetic process
Chronic inflammatory response
Response to peptide
PID HIF1 TFPATHWAY
PID TOLL ENDOGENOUS PATHWAY
Regulation of hormone biosynthetic process
Antigen processing and presentation
Interactions of Vpr with host cellular proteins
Nop56p-associated pre-rRNA complex
Myeloid leukocyte activation
Cofactor catabolic process
Adaptive immune response
Viral life cycle
NABA ECM REGULATORS
IL1B_KC
Response to unfolded protein
Signaling by Interleukins
Antigen processing and presentation
Regulation of innate immune response
Interferon-gamma-mediated signaling pathway
Positive regulation of protein kinase activity
Reactive oxygen species metabolic process
Response to inorganic substance
Negative regulation of intracellular signal transduction
Lymphocyte activation
TNF-alpha/Nf-kappa B signaling complex
Regulation of transforming growth factor beta receptor signaling pathway
Response to nutrient levels
Alpha-beta T cell activation
Response to oxygen levels
Establishment of protein localization to organelle
Response to bacterium
Response to radiation
Response to hexose
Response to toxic substance
Positive regulation of cell activation
Response to antibiotic
Leukocyte activation involved in immune response
Response to inorganic substance
Tuberculosis
Cell chemotaxis
Regulation of ion transport
Regulation of actin cytoskeleton organization
Toll-like receptor 4 signaling pathway
Signaling by RAF1 mutants
Immune response-activating signal transduction
Negative regulation of reproductive process
Membrane fusion
ERK1 and ERK2 cascade
Transcriptional misregulation in cancer
Positive regulation of response to external stimulus
Regulation of metal ion transport
PID CXCR4 PATHWAY
Regulation of secretion by cell
Response to steroid hormone
-log10(P)

### Figure S4

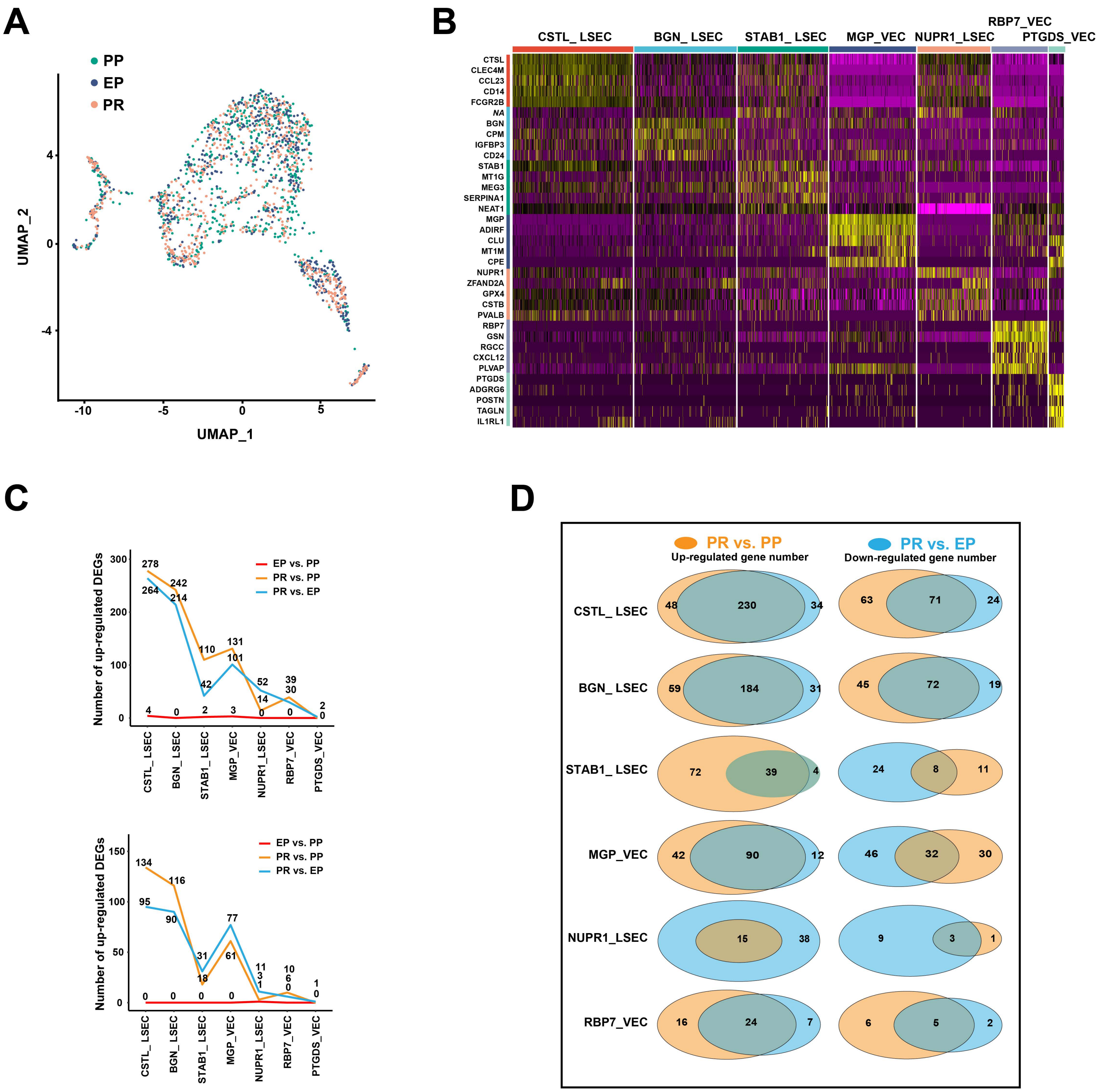

A
PP
EP
PR
UMAP_2
UMAP_1
B
CSTL_ LSEC
BGN_ LSEC
STAB1_ LSEC
MGP_VEC
NUPR1_LSEC
RBP7_VEC
PTGDS_VEC
CTSL
CLEC4M
CCL23
CD14
FCGR2B
NA
BGN
CPM
IGFBP3
CD24
STAB1
MT1G
MEG3
SERPINA1
NEAT1
MGP
ADIRF
CLU
MT1M
CPE
NUPR1
ZFAND2A
GPX4
CSTB
PVALB
RBP7
GSN
RGCC
CXCL12
PLVAP
PTGDS
ADGRG6
POSTN
TAGLN
IL1RL1
C
Number of up-regulated DEGs
EP vs. PP
PR vs. PP
PR vs. EP
CSTL_LSEC
BGN_LSEC
STAB1_LSEC
MGP_VEC
NUPR1_LSEC
RBP7_VEC
PTGDS_VEC
D
PR vs. PP
PR vs. EP
Up-regulated gene number
Down-regulated gene number
CSTL_ LSEC
BGN_ LSEC
STAB1_ LSEC
MGP_VEC
NUPR1_LSEC
RBP7_VEC

### Figure S5

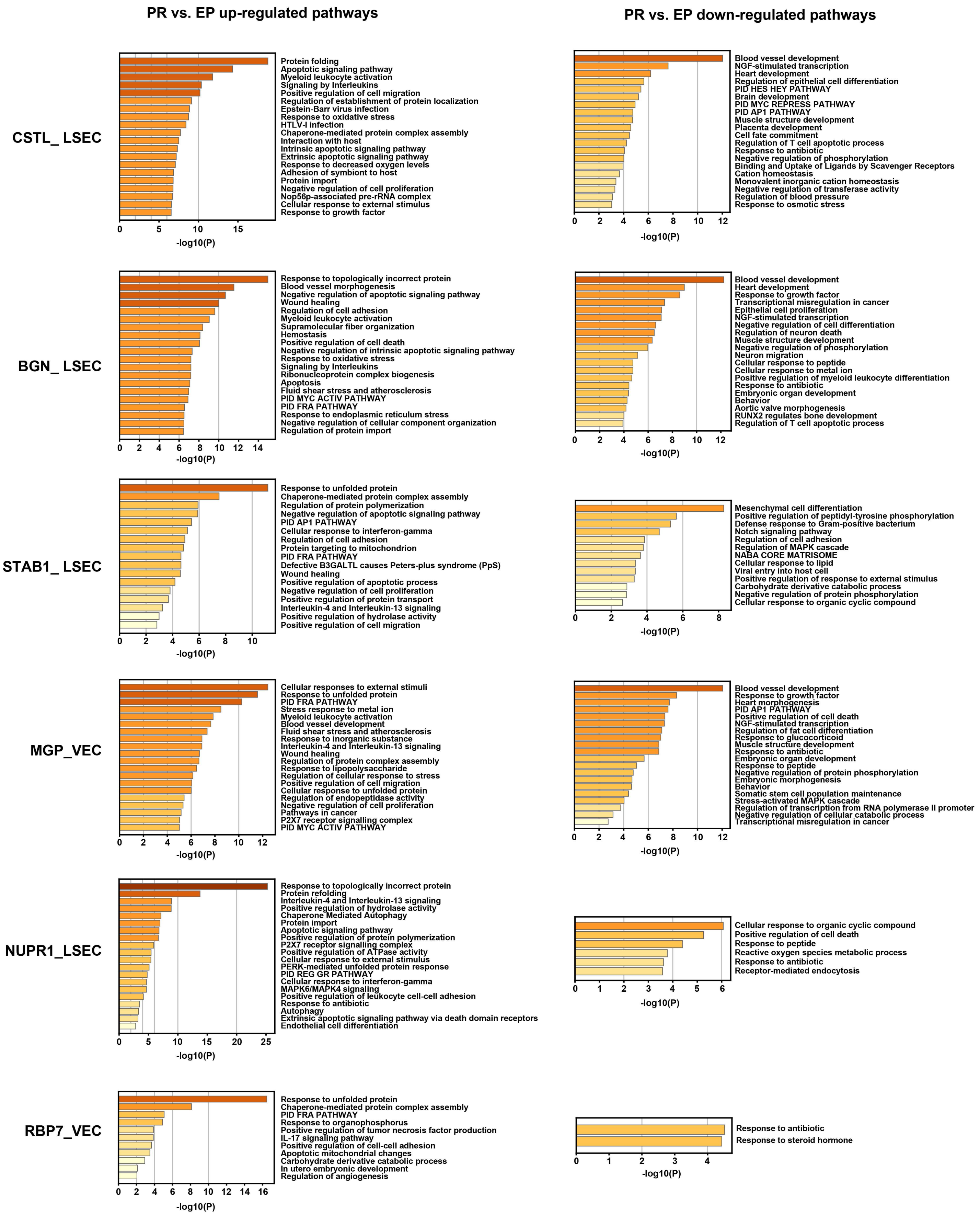
PR vs. EP up-regulated pathways
PR vs. EP down-regulated pathways
CSTL_ LSEC
Protein folding
Apoptotic signaling pathway
Myeloid leukocyte activation
Signaling by Interleukins
Positive regulation of cell migration
Regulation of establishment of protein localization
Epstein-Barr virus infection
Response to oxidative stress
HTLV-I infection
Chaperone-mediated protein complex assembly
Interaction with host
Intrinsic apoptotic signaling pathway
Extrinsic apoptotic signaling pathway
Response to decreased oxygen levels
Adhesion of symbiont to host
Protein import
Negative regulation of cell proliferation
Nop56p-associated pre-rRNA complex
Cellular response to external stimulus
Response to growth factor
0 5 10 15
-log10(P)
Blood vessel development
NGF-stimulated transcription
Heart development
Regulation of epithelial cell differentiation
PID HES HEY PATHWAY
Brain development
PID MYC REPRESS PATHWAY
PID AP1 PATHWAY
Muscle structure development
Placenta development
Cell fate commitment
Regulation of T cell apoptotic process
Response to antibiotic
Negative regulation of phosphorylation
Binding and Uptake of Ligands by Scavenger Receptors
Cation homeostasis
Monovalent inorganic cation homeostasis
Negative regulation of transferase activity
Regulation of blood pressure
Response to osmotic stress
0 2 4 6 8 10 12
-log10(P)
BGN_ LSEC
Response to topologically incorrect protein
Blood vessel morphogenesis
Negative regulation of apoptotic signaling pathway
Wound healing
Regulation of cell adhesion
Myeloid leukocyte activation
Supramolecular fiber organization
Hemostasis
Positive regulation of cell death
Negative regulation of intrinsic apoptotic signaling pathway
Response to oxidative stress
Signaling by Interleukins
Ribonucleoprotein complex biogenesis
Apoptosis
Fluid shear stress and atherosclerosis
PID MYC ACTIV PATHWAY
PID FRA PATHWAY
Response to endoplasmic reticulum stress
Negative regulation of cellular component organization
Regulation of protein import
0 2 4 6 8 10 12 14
-log10(P)
Blood vessel development
Heart development
Response to growth factor
Transcriptional misregulation in cancer
Epithelial cell proliferation
NGF-stimulated transcription
Negative regulation of cell differentiation
Regulation of neuron death
Muscle structure development
Negative regulation of phosphorylation
Neuron migration
Cellular response to peptide
Cellular response to metal ion
Positive regulation of myeloid leukocyte differentiation
Response to antibiotic
Embryonic organ development
Behavior
Aortic valve morphogenesis
RUNX2 regulates bone development
Regulation of T cell apoptotic process
0 2 4 6 8 10 12
-log10(P)
STAB1_ LSEC
Response to unfolded protein
Chaperone-mediated protein complex assembly
Regulation of protein polymerization
Negative regulation of apoptotic signaling pathway
PID AP1 PATHWAY
Cellular response to interferon-gamma
Regulation of cell adhesion
Protein targeting to mitochondrion
PID FRA PATHWAY
Defective B3GALTL causes Peters-plus syndrome (PpS)
Wound healing
Positive regulation of apoptotic process
Negative regulation of cell proliferation
Positive regulation of protein transport
Interleukin-4 and Interleukin-13 signaling
Positive regulation of hydrolase activity
Positive regulation of cell migration
0 2 4 6 8 10
-log10(P)
Mesenchymal cell differentiation
Positive regulation of peptidyl-tyrosine phosphorylation
Defense response to Gram-positive bacterium
Notch signaling pathway
Regulation of cell adhesion
Regulation of MAPK cascade
NABA CORE MATRISOME
Cellular response to lipid
Viral entry into host cell
Positive regulation of response to external stimulus
Carbohydrate derivative catabolic process
Negative regulation of protein phosphorylation
Cellular response to organic cyclic compound
0 2 4 6 8
-log10(P)
MGP_VEC
Cellular responses to external stimuli
Response to unfolded protein
PID FRA PATHWAY
Stress response to metal ion
Myeloid leukocyte activation
Blood vessel development
Fluid shear stress and atherosclerosis
Response to inorganic substance
Interleukin-4 and Interleukin-13 signaling
Wound healing
Regulation of protein complex assembly
Response to lipopolysaccharide
Regulation of cellular response to stress
Positive regulation of cell migration
Cellular response to unfolded protein
Regulation of endopeptidase activity
Negative regulation of cell proliferation
Pathways in cancer
P2X7 receptor signalling complex
PID MYC ACTIV PATHWAY
0 2 4 6 8 10 12
-log10(P)
Blood vessel development
Response to growth factor
Heart morphogenesis
PID AP1 PATHWAY
Positive regulation of cell death
NGF-stimulated transcription
Regulation of fat cell differentiation
Response to glucocorticoid
Muscle structure development
Response to antibiotic
Embryonic organ development
Response to peptide
Negative regulation of protein phosphorylation
Embryonic morphogenesis
Behavior
Somatic stem cell population maintenance
Stress-activated MAPK cascade
Regulation of transcription from RNA polymerase II promoter
Negative regulation of cellular catabolic process
Transcriptional misregulation in cancer
0 2 4 6 8 10 12
-log10(P)
NUPR1_LSEC
Response to topologically incorrect protein
Protein refolding
Interleukin-4 and Interleukin-13 signaling
Positive regulation of hydrolase activity
Chaperone Mediated Autophagy
Protein import
Apoptotic signaling pathway
Positive regulation of protein polymerization
P2X7 receptor signalling complex
Positive regulation of ATPase activity
Cellular response to external stimulus
PERK-mediated unfolded protein response
PID REG GR PATHWAY
Cellular response to interferon-gamma
MAPK6/MAPK4 signaling
Positive regulation of leukocyte cell-cell adhesion
Response to antibiotic
Autophagy
Extrinsic apoptotic signaling pathway via death domain receptors
Endothelial cell differentiation
0 5 10 15 20 25
-log10(P)
Cellular response to organic cyclic compound
Positive regulation of cell death
Response to peptide
Reactive oxygen species metabolic process
Response to antibiotic
Receptor-mediated endocytosis
0 1 2 3 4 5 6
-log10(P)
RBP7_VEC
Response to unfolded protein
Chaperone-mediated protein complex assembly
PID FRA PATHWAY
Response to organophosphorus
Positive regulation of tumor necrosis factor production
IL-17 signaling pathway
Positive regulation of cell-cell adhesion
Apoptotic mitochondrial changes
Carbohydrate derivative catabolic process
In utero embryonic development
Regulation of angiogenesis
0 2 4 6 8 10 12 14 16
-log10(P)
Response to antibiotic
Response to steroid hormone
0 1 2 3 4
-log10(P)

### Figure S6

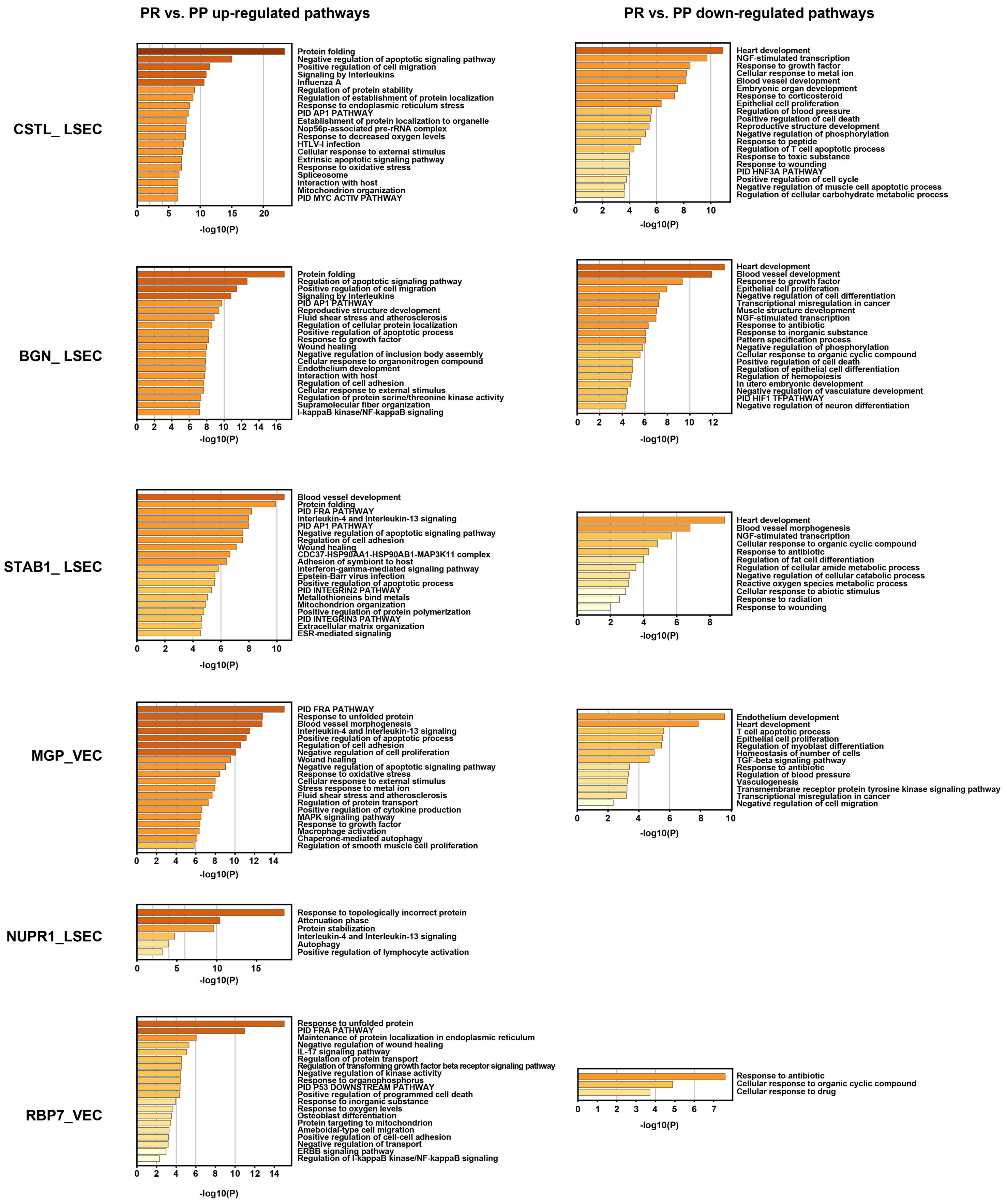
PR vs. PP up-regulated pathways
PR vs. PP down-regulated pathways
CSTL_ LSEC
Protein folding
Negative regulation of apoptotic signaling pathway
Positive regulation of cell migration
Signaling by Interleukins
Influenza A
Regulation of protein stability
Regulation of establishment of protein localization
Response to endoplasmic reticulum stress
PID AP1 PATHWAY
Establishment of protein localization to organelle
Nop56p-associated pre-rRNA complex
Response to decreased oxygen levels
HTLV-I infection
Cellular response to external stimulus
Extrinsic apoptotic signaling pathway
Response to oxidative stress
Spliceosome
Interaction with host
Mitochondrion organization
PID MYC ACTIV PATHWAY
-log10(P)
Heart development
NGF-stimulated transcription
Response to growth factor
Cellular response to metal ion
Blood vessel development
Embryonic organ development
Response to corticosteroid
Epithelial cell proliferation
Regulation of blood pressure
Positive regulation of cell death
Reproductive structure development
Negative regulation of phosphorylation
Response to peptide
Regulation of T cell apoptotic process
Response to toxic substance
Response to wounding
PID HNF3A PATHWAY
Positive regulation of cell cycle
Negative regulation of muscle cell apoptotic process
Regulation of cellular carbohydrate metabolic process
-log10(P)
BGN_ LSEC
Protein folding
Regulation of apoptotic signaling pathway
Positive regulation of cell migration
Signaling by Interleukins
PID AP1 PATHWAY
Reproductive structure development
Fluid shear stress and atherosclerosis
Regulation of cellular protein localization
Positive regulation of apoptotic process
Response to growth factor
Wound healing
Negative regulation of inclusion body assembly
Cellular response to organonitrogen compound
Endothelium development
Interaction with host
Regulation of cell adhesion
Cellular response to external stimulus
Regulation of protein serine/threonine kinase activity
Supramolecular fiber organization
I-kappaB kinase/NF-kappaB signaling
-log10(P)
Heart development
Blood vessel development
Response to growth factor
Epithelial cell proliferation
Negative regulation of cell differentiation
Transcriptional misregulation in cancer
Muscle structure development
NGF-stimulated transcription
Response to antibiotic
Response to inorganic substance
Pattern specification process
Negative regulation of phosphorylation
Cellular response to organic cyclic compound
Positive regulation of cell death
Regulation of epithelial cell differentiation
Regulation of hemopoiesis
In utero embryonic development
Negative regulation of vasculature development
PID HIF1 TFPATHWAY
Negative regulation of neuron differentiation
-log10(P)
STAB1_ LSEC
Blood vessel development
Protein folding
PID FRA PATHWAY
Interleukin-4 and Interleukin-13 signaling
PID AP1 PATHWAY
Negative regulation of apoptotic signaling pathway
Regulation of cell adhesion
Wound healing
CDC37-HSP90AA1-HSP90AB1-MAP3K11 complex
Adhesion of symbiont to host
Interferon-gamma-mediated signaling pathway
Epstein-Barr virus infection
Positive regulation of apoptotic process
PID INTEGRIN2 PATHWAY
Metallothioneins bind metals
Mitochondrion organization
Positive regulation of protein polymerization
PID INTEGRIN3 PATHWAY
Extracellular matrix organization
ESR-mediated signaling
-log10(P)
Heart development
Blood vessel morphogenesis
NGF-stimulated transcription
Cellular response to organic cyclic compound
Response to antibiotic
Regulation of fat cell differentiation
Regulation of cellular amide metabolic process
Negative regulation of cellular catabolic process
Reactive oxygen species metabolic process
Cellular response to abiotic stimulus
Response to radiation
Response to wounding
-log10(P)
MGP_VEC
PID FRA PATHWAY
Response to unfolded protein
Blood vessel morphogenesis
Interleukin-4 and Interleukin-13 signaling
Positive regulation of apoptotic process
Regulation of cell adhesion
Negative regulation of cell proliferation
Wound healing
Negative regulation of apoptotic signaling pathway
Response to oxidative stress
Cellular response to external stimulus
Stress response to metal ion
Fluid shear stress and atherosclerosis
Regulation of protein transport
Positive regulation of cytokine production
MAPK signaling pathway
Response to growth factor
Macrophage activation
Chaperone-mediated autophagy
Regulation of smooth muscle cell proliferation
-log10(P)
Endothelium development
Heart development
T cell apoptotic process
Epithelial cell proliferation
Regulation of myoblast differentiation
Homeostasis of number of cells
TGF-beta signaling pathway
Response to antibiotic
Regulation of blood pressure
Vasculogenesis
Transmembrane receptor protein tyrosine kinase signaling pathway
Transcriptional misregulation in cancer
Negative regulation of cell migration
-log10(P)
NUPR1_LSEC
Response to topologically incorrect protein
Attenuation phase
Protein stabilization
Interleukin-4 and Interleukin-13 signaling
Autophagy
Positive regulation of lymphocyte activation
-log10(P)
RBP7_VEC
Response to unfolded protein
PID FRA PATHWAY
Maintenance of protein localization in endoplasmic reticulum
Negative regulation of wound healing
IL-17 signaling pathway
Regulation of protein transport
Regulation of transforming growth factor beta receptor signaling pathway
Negative regulation of kinase activity
Response to organophosphorus
PID P53 DOWNSTREAM PATHWAY
Positive regulation of programmed cell death
Response to inorganic substance
Response to oxygen levels
Osteoblast differentiation
Protein targeting to mitochondrion
Ameboidal-type cell migration
Positive regulation of cell-cell adhesion
Negative regulation of transport
ERBB signaling pathway
Regulation of I-kappaB kinase/NF-kappaB signaling
-log10(P)
Response to antibiotic
Cellular response to organic cyclic compound
Cellular response to drug
-log10(P)

**Figure S7**

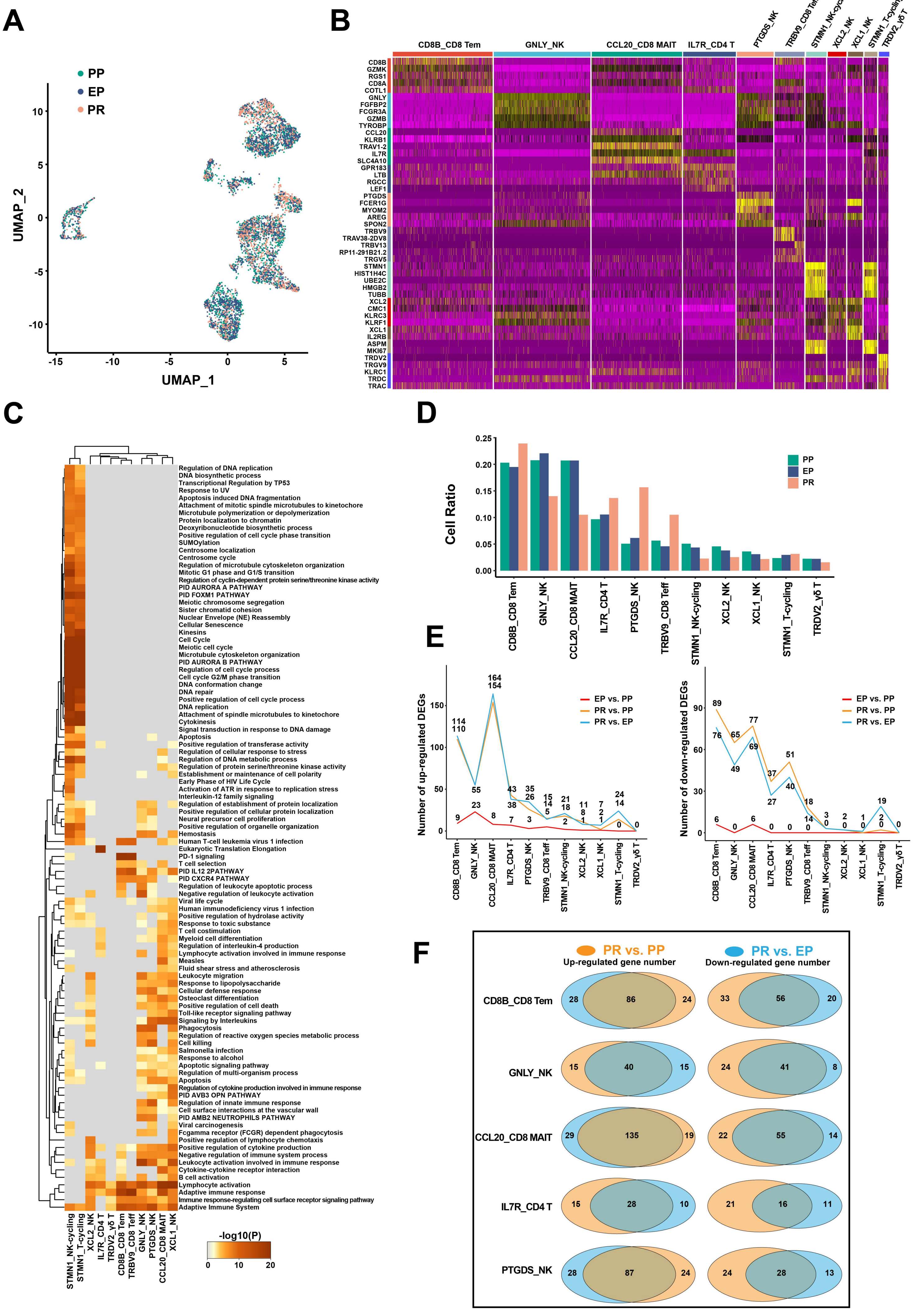
A
PP
EP
PR
UMAP_1
UMAP_2
B
CD8B_CD8 Tem
GNLY_NK
CCL20_CD8 MAIT
IL7R_CD4 T
PTGDS_NK
TRBV9_CD8 Teff
STMN1_NK-cycling
XCL2_NK
XCL1_NK
STMN1_T-cycling
TRDV2_γδ T
C
-log10(P)
D
Cell Ratio
E
Number of up-regulated DEGs
Number of down-regulated DEGs
EP vs. PP
PR vs. PP
PR vs. EP
F
PR vs. PP
Up-regulated gene number
PR vs. EP
Down-regulated gene number

### Figure S8

**A**

PR vs. EP up-regulated pathways

PR vs. EP down-regulated pathways

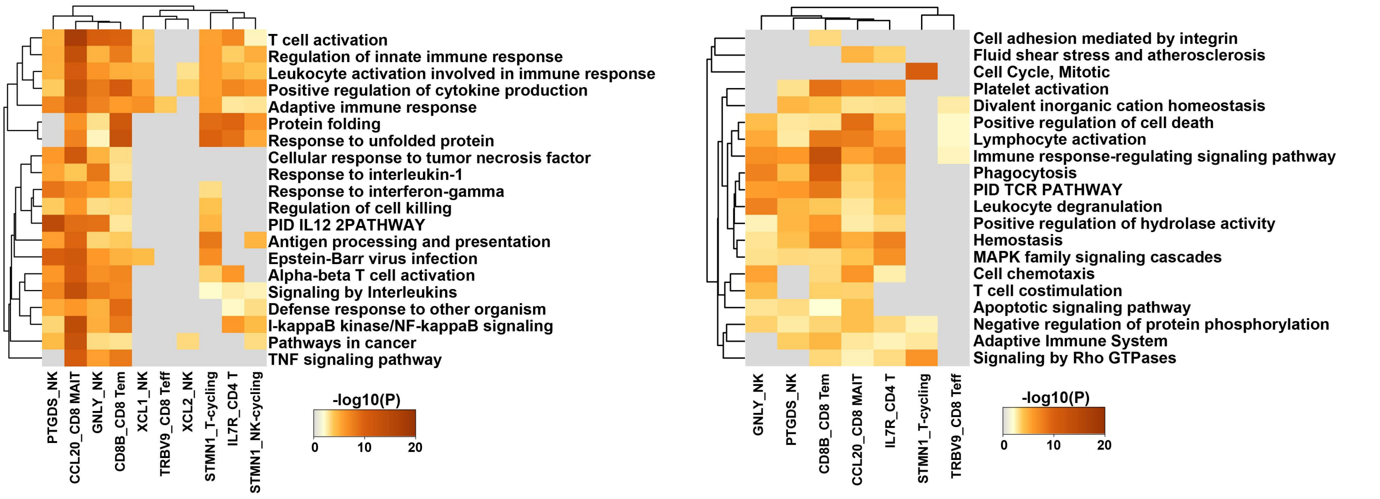


**B**

PR vs. PP up-regulated pathways

PR vs. PP down-regulated pathways

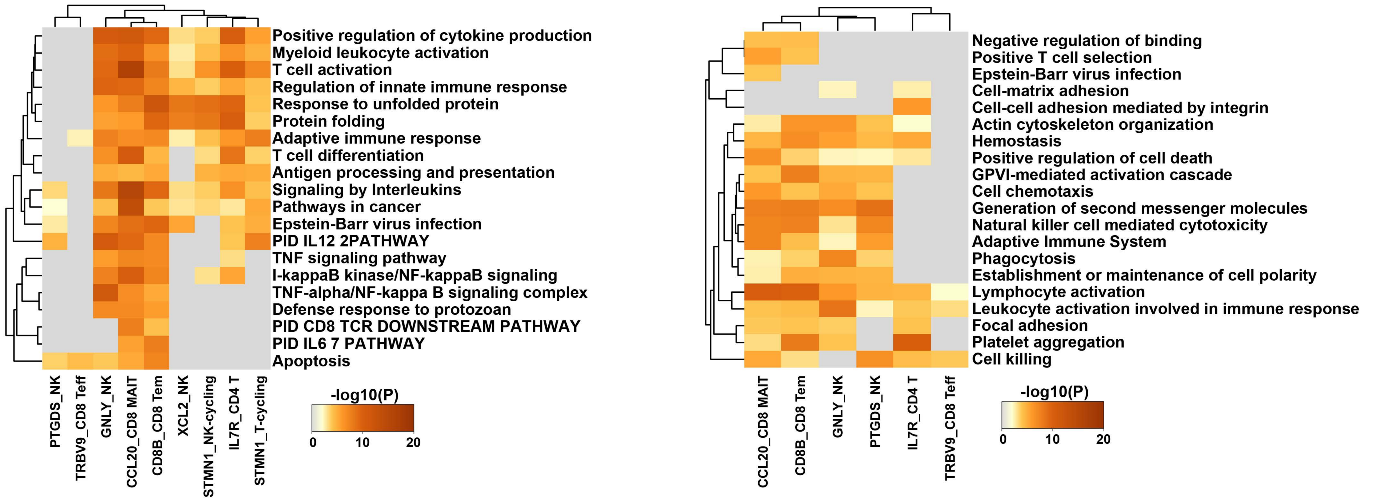

### Figure S9

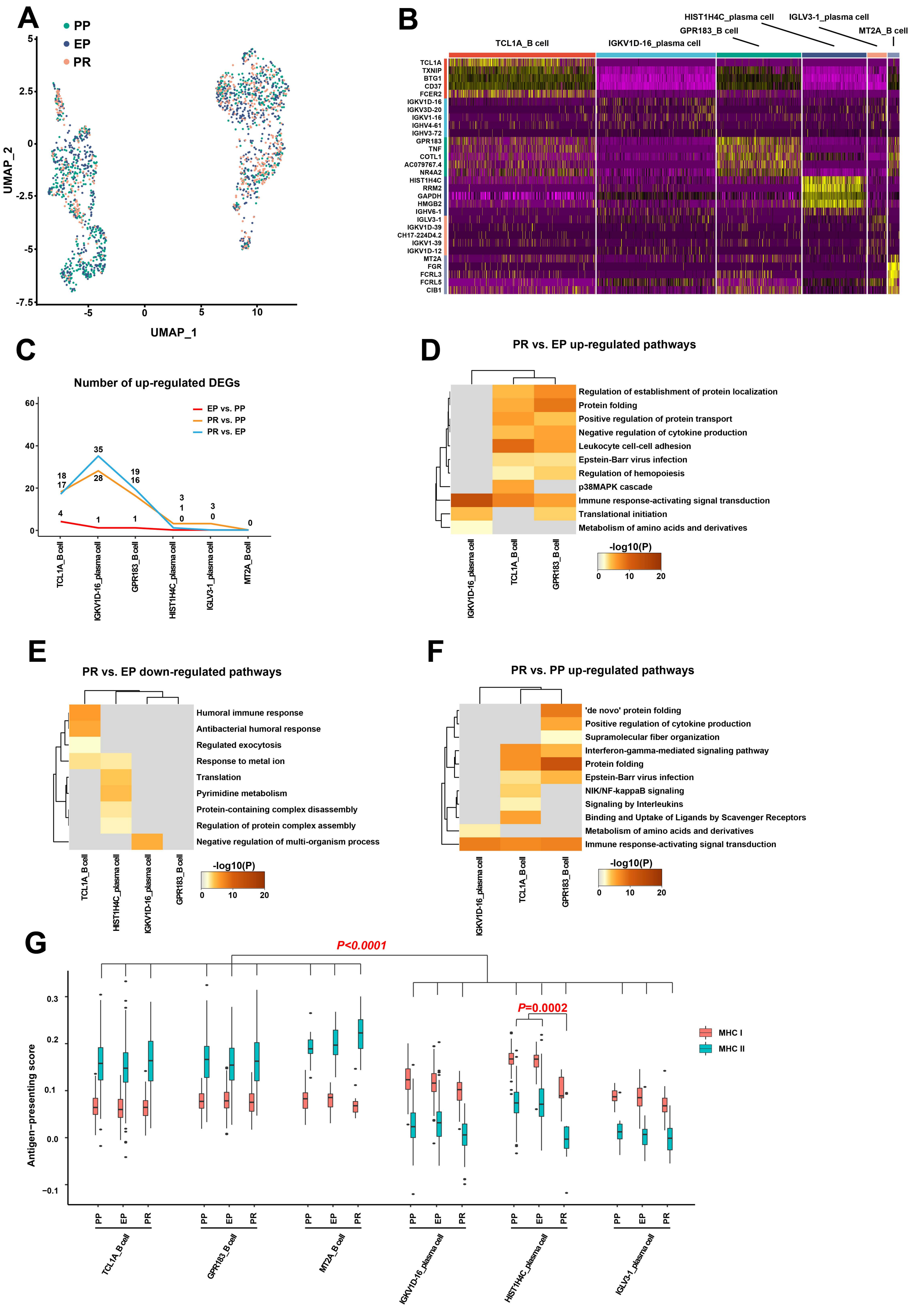

A
PP
EP
PR
UMAP_1
UMAP_2
B
TCL1A_B cell
IGKV1D-16_plasma cell
GPR183_B cell
HIST1H4C_plasma cell
IGLV3-1_plasma cell
MT2A_B cell
TCL1A
TXNIP
BTG1
CD37
FCER2
IGKV1D-16
IGKV3D-20
IGKV1-16
IGHV4-61
IGHV3-72
GPR183
TNF
COTL1
AC079767.4
NR4A2
HIST1H4C
RRM2
GAPDH
HMGB2
IGHV6-1
IGLV3-1
IGKV1D-39
CH17-224D4.2
IGKV1-39
IGKV1D-12
MT2A
FGR
FCRL3
FCRL5
CIB1
C
Number of up-regulated DEGs
EP vs. PP
PR vs. PP
PR vs. EP
D
PR vs. EP up-regulated pathways
Regulation of establishment of protein localization
Protein folding
Positive regulation of protein transport
Negative regulation of cytokine production
Leukocyte cell-cell adhesion
Epstein-Barr virus infection
Regulation of hemopoiesis
p38MAPK cascade
Immune response-activating signal transduction
Translational initiation
Metabolism of amino acids and derivatives
-log10(P)
E
PR vs. EP down-regulated pathways
Humoral immune response
Antibacterial humoral response
Regulated exocytosis
Response to metal ion
Translation
Pyrimidine metabolism
Protein-containing complex disassembly
Regulation of protein complex assembly
Negative regulation of multi-organism process
-log10(P)
F
PR vs. PP up-regulated pathways
'de novo' protein folding
Positive regulation of cytokine production
Supramolecular fiber organization
Interferon-gamma-mediated signaling pathway
Protein folding
Epstein-Barr virus infection
NIK/NF-kappaB signaling
Signaling by Interleukins
Binding and Uptake of Ligands by Scavenger Receptors
Metabolism of amino acids and derivatives
Immune response-activating signal transduction
-log10(P)
G
P<0.0001
P=0.0002
Antigen−presenting score
MHC I
MHC II

### Figure Legend

**Figure S1:**

**A**: “DoubletFinder” software identified 1160 doublet cells in PP, EP and PR. Red dots represent high expression level doublets, green dots represent low level doublets, and blue dots represent singlet.

**B**: Feature plots showing the normalized expression of marker genes in different cell lineage.

**C**: UMAP visualization of all cell clusters by PP, EP ad PR samples.

**D**: Violin plots showing the normalized expression of *MARCO* and *CD68* genes (y axis) for mononuclear phagocyte clusters (x axis).

**Figure S2:**

**A**: UMAP visualization of mononuclear phagocyte clusters by PP, EP ad PR samples.

**B**: Gene Ontology enrichment analysis results of mononuclear phagocyte clusters. The top 100 most significant GO terms ($P$-value < 0.05) are shown in rows.

**C**: Number of down-regulated DEGs between different samples in different mononuclear phagocyte clusters.

**D**: The Venn diagram shows the overlap gene number of DEGs in PR vs. PP (yellow) and DEGs in PR vs. EP (blue) of different mononuclear phagocyte clusters. The left column shows the number of up-regulated genes while the right column shows the number of down-regulated genes.

**E**: Antigen presentation ability of MHC class I genes (red) and MHC class II genes (green) in different mononuclear macrophage clusters. The antigen-presentating score (y axis) of MHC class II genes of KC clusters is higher than that of TMo clusters (x axis, $P$<0.0001).

**Figure S3:**

**A-B**: Gene Ontology enrichment analysis results of up-regulated (left column) and down-regulated DEGs (right column) in different mononuclear phagocyte clusters from PR vs. EP (**Figure S3A**) and PR vs. PP (**Figure S3B**). Only the top 20 most significant GO terms ($P$-value < 0.05) are shown in rows.

**Figure S4:**

**A**: UMAP visualization of endothelial cell clusters by PP, EP ad PR samples.

**B**: Heatmap of top five differentially expressed genes between different endothelial cell clusters. The line is colored according to clusters in **Figure 4A**.

**C**: Number of up-regulated (left panel) and up-regulated (right panel) DEGs between different samples in different endothelial cell clusters.

**D**: The Venn diagram shows the overlap gene number of DEGs in PR vs. PP (yellow) and DEGs in PR vs. EP (blue) of different endothelial cell clusters. The left column shows the number of up-regulated genes while the right column shows the number of down-regulated genes.

**Figure S5:**

Gene Ontology enrichment analysis results of up-regulated (left column) and down-regulated DEGs (right column) in different endothelial cell clusters from PR vs. EP. Only the top 20 most significant GO terms ($P$-value < 0.05) are shown in rows.

**Figure S6:**

Gene Ontology enrichment analysis results of up-regulated (left column) and down-regulated DEGs (right column) in different endothelial cell clusters from PR vs. PP. Only the top 20 most significant GO terms ($P$-value < 0.05) are shown in rows.

**Figure S7:**

**A**: UMAP visualization of NK/T cell clusters by PP, EP ad PR samples.

**B**: Heatmap of top five differentially expressed genes between different NK/T cell clusters. The line is colored according to clusters in **Figure 5A**.

**C**: Gene Ontology enrichment analysis results of NK/T cell clusters. Only the top 100 most significant GO terms (*P*-value < 0.05) are shown in rows.

**D**: Cell ratio of different NK/T cell clusters in PP, EP and PR samples.

**E**: Number of up-regulated (left panel) and up-regulated (right panel) DEGs between different samples in different NK/T cell clusters.

**F**: The Venn diagram shows the overlap gene number of DEGs in PR vs. PP (yellow) and DEGs in PR vs. EP (blue) of different NK/T cell clusters. The left column shows the number of up-regulated genes while the right column shows the number of down-regulated genes.

**Figure S8:**

**A-B**: Gene Ontology enrichment analysis results of up-regulated (left panel) and down-regulated DEGs (right panel) in different mononuclear phagocyte clusters from PR vs. EP (**Figure S8A**) and PR vs. PP (**Figure S8B**). Only the top 20 most significant GO terms (*P*-value < 0.05) are shown in rows.

**Figure S9:**

**A**: UMAP visualization of B and plasma cell clusters by PP, EP ad PR samples.

**B**: Heatmap of top five differentially expressed genes between different B and plasma cell clusters. The line is colored according to clusters in **Figure 6A**.

**C**: Number of up-regulated DEGs between different samples in different B and plasma cell clusters.

**D-E**: Gene Ontology enrichment analysis results of up-regulated pathways (**Figure S9D**) and down-regulated (**Figure S9E**) in different B and plasma cell clusters from PR vs. EP.

**F**: Gene Ontology enrichment analysis results of up-regulated pathways in different B and plasma cell clusters from PR vs. PP.

**G**: Antigen presentation ability of MHC class I genes (red) and MHC class II genes (green) in different B and plasma cell clusters. The antigen-presentating score (y axis) of MHC class II genes of B cell clusters is higher than that of plasma cell clusters (x axis, *P*<0.0001), and decrease after reperfusion in HIST1H4C_plasma (*P*=0.0002).
